# Endothelial KRAS^G12V^ signaling drives aberrant morphogenesis and establishes an AVM transcriptional identity in primary human endothelial cells

**DOI:** 10.64898/2026.06.26.734855

**Authors:** Samantha King, Qing-fen Li, Ramon Bossardi Ramos, Kevin Pumiglia

**Affiliations:** Department of Regenerative and Cancer Cell Biology at Albany Medical College; Department of Molecular and Cellular Physiology, Albany Medical College, 47 New Scotland Avenue, Albany, NY 12208

**Author notes:** Kevin Pumiglia |.

**Keywords:** brain arteriovenous malformation, KRAS, endothelial cell, PI3K, TRAP-seq, AVM transcriptional identity

## Abstract

Somatic activating mutations in KRAS are found in the endothelium of the majority of sporadic brain arteriovenous malformations (bAVMs), yet the consequences of oncogenic KRAS signaling in endothelial cells during active vessel morphogenesis remain incompletely characterized. We expressed KRAS^G12V^ in primary human umbilical vein endothelial cells using a doxycycline-inducible lentiviral system and examined morphogenic behavior, proliferation, migration, and transcriptional output in a three-dimensional planar co-culture angiogenesis assay. KRAS^G12V^-expressing cells failed to organize into vessel-like networks, instead forming compact sheet-like structures that persisted through day 12. A transient proliferative phase at days 3–5 resolved to control levels by day 12, consistent with preserved sensitivity to contact inhibition rather than unrestricted growth. Enhanced migration at day 5 was accompanied by upregulation of a focal adhesion and matrix remodeling program centered on ITGB3, PLAU, PLAUR, and PIK3CG. Translating ribosome-affinity purification sequencing (TRAP-seq) of the EC-specific translatome across four independent donor pools revealed progressive acquisition of an AVM-associated transcriptional identity by day 12, including upregulation of ACVRL1, ENG, JAG1, NOTCH1, ANGPT2, and TEK, with concordance to human bAVM nidus endothelium at both the gene and pathway level. Pharmacological inhibition with Alpelisib (PI3Kα), Trametinib (MEK), and Pazopanib (VEGFR2) demonstrated that PI3K is the principal organizer of the morphogenic phenotype. These findings characterize a KRASG12V-driven program in endothelial cells that recapitulates core transcriptional features of bAVM endothelium in a primary cell model.

## INTRODUCTION

Arteriovenous malformations are structurally abnormal vascular lesions in which arteries and veins form direct connections that bypass the intervening capillary bed. The resulting high-pressure shunts are prone to rupture, and brain AVMs represent a leading cause of hemorrhagic stroke in children and young adults [1]. Current management relies on surgical resection, stereotactic radiosurgery, or endovascular embolization, as no pharmacological therapy is approved specifically for AVM. This gap in treatment reflects, in part, an incomplete understanding of the cell-intrinsic events that initiate malformation.

A significant advance in this area came from the discovery that somatic activating KRAS mutations are present in the endothelial cells of the majority of sporadic brain AVMs, with KRAS^G12V^ and KRAS^G12D^ among the most identified variants [2]. In endothelial cell culture studies derived from that work, mutant KRAS increased ERK activity, enhanced migration, and upregulated genes associated with angiogenesis and Notch signaling, effects attributable to MAPK/ERK pathway activation [2]. The identification of somatic KRAS mutations in sporadic AVMs has also provided a point of convergence with hereditary vascular malformation syndromes. Hereditary hemorrhagic telangiectasia arises from germline loss-of-function mutations in the BMP/TGF-β pathway components ACVRL1 (ALK1) and ENG (Endoglin), and several lines of evidence indicate that activation of the PI3K/mTOR pathway in ECs contributes to its pathobiology. EC proliferation in cutaneous telangiectasias from HHT1 patients is associated with elevated PI3K pathway markers, including pS6 and pNDRG1 [3], and combined correction of mTOR and VEGFR2 activity reversed retinal AVMs and reduced gastrointestinal bleeding in HHT mouse models [4]. That PI3K/mTOR involvement in AVM pathobiology extends across genetically distinct subtypes raises the question of whether PI3K signaling also contributes to KRAS-driven malformation in the endothelium.[2-4]

In vivo models of endothelial KRAS^G12V^ expression have established that the mutation is sufficient to induce vascular malformations and that ERK activation is a consistent feature, but they have not fully resolved the mechanisms driving the phenotype. Fish et al. used endothelial-specific KRAS^G12D^ delivery in mice and KRAS^G12D^ or KRAS^G12V^ in zebrafish (with no substantive phenotypic differences observed between effector mutants) to produce AVMs characterized principally by increased EC size and abnormal migration, with no evidence of frank EC proliferation and no requirement for PI3K signaling, as MEK inhibition was sufficient to suppress malformation in that system [5]. Park et al. generated brain AVMs in mice using an AAV vector and reported focal angiogenesis associated with KRAS^G12V^ expression, with lesion growth suppressed by trametinib [6]. Saito et al. similarly induced postnatal brain AVMs via AAV and found occasional Ki67 signal in lesion ECs but concluded that cellular growth and dilation were the predominant contributors rather than proliferation [7]. In contrast, endothelial HRAS^V12^ expression in the mouse produces cerebrovascular malformations accompanied by EC proliferation in vivo [8]. This suggests that the proliferative response to activated Ras in the endothelium may depend on the specific isoform and the vascular context in which it is expressed, such as the organ-specific transcriptional identity of the endothelium, the developmental stage of expression, and/or the proportion of cells carrying the mutation. Consistent with this, KRAS^G12D^ in primary human lymphatic endothelial cells is sufficient to induce proliferation and migration in culture [9] and prior work from our laboratory and others has established activated Ras as sufficient to drive EC proliferation and survival in vitro [10-13]. These data support a proliferative capacity for activated Ras in endothelial cells whose in vivo expression appears context dependent. Together, the variability across models in the extent of EC proliferation and the pathway dependencies identified in different systems suggest that the effects of KRAS^G12V^ signaling in the endothelium during active morphogenesis remain unresolved.[5-13]

PI3K signaling, particularly through the p110α catalytic isoform, is a well-established requirement for EC migration during angiogenesis [14], and PI3K-dependent inhibition of actomyosin contractility is required for the EC rearrangements that underlie vascular patterning [15]. KRAS signals through both the MAPK/ERK and PI3K/AKT pathways, and the relative engagement of these pathways varies with cellular context and the intensity of upstream activation [16]. Whether KRAS^G12V^ engages PI3K-dependent EC behaviors during active morphogenesis, and whether this can be separated from MAPK-dependent effects, requires a system in which ECs undergo morphogenesis under controlled pharmacological and genetic conditions.

The 3D planar co-culture assay, in which primary endothelial cells organize into vessel-like structures over a fibroblast monolayer, provides a well-characterized model of EC morphogenesis that recapitulates progressive tubulogenesis, cell-cell contact, and quiescence over a 12-day period [17]. Because ECs constitute a minority of the total cell population in this mixed-culture system, conventional bulk RNA-seq cannot resolve their transcriptional state from that of the fibroblast background. We recently described the application of TRAP-seq using a lentiviral HA-Rpl22 ribosome-tagging approach selectively expressed in HUVECs, which permits the isolation of actively translating EC mRNA from co-cultures across the morphogenetic time course [18]. That study demonstrated that TRAP-seq selectively enriches endothelial transcripts and captures the temporal dynamics of normal morphogenesis, with blood vessel development, cell cycle regulation, and Notch pathway genes among the most informative outputs. Because KRAS^G12V^ in the present study is delivered to HUVECs by a doxycycline-inducible lentiviral vector, its expression is restricted to the EC compartment while the fibroblast microenvironment, including secreted matrix, paracrine signals, and physical support, is preserved. This combination of cell-type-specific transcriptomics and an EC-restricted genetic perturbation provides resolution that neither bulk sequencing nor conventional 2D culture can offer.

Here we report that KRAS^G12V^ expression in primary HUVECs in 3D co-culture produces a coherent program of morphological disruption, enhanced migration, and progressive acquisition of an AVM/arterial transcriptional identity that includes upregulation of Notch pathway components and genes associated with arteriovenous identity. A transient proliferative phase accompanies early morphogenesis but resolves with contact, consistent with preserved sensitivity to contact inhibition rather than a fundamental override of quiescence. PI3K inhibition with Alpelisib selectively rescued all three phenotypic components and restored near-normal vessel architecture, while the MEK inhibitor Trametinib and the VEGFR2 inhibitor Pazopanib were only partially effective, identifying PI3K as the principal driver of the defective morphogenesis in this system.

## METHODS

### Cell Culture

Human umbilical vein endothelial cells (HUVECs) from pooled donors were purchased from Lonza and PromoCell and cultured as we have previously described [18]. HUVECs were maintained in endothelial complete growth medium-2 (ECGM2, PromoCell) and used at low passage (P1-P7). Cells were maintained in the absence of doxycycline prior to experimental induction. Primary normal human dermal fibroblasts (NHDFs) were purchased from PromoCell and maintained in DMEM supplemented with 10% FBS.

### Plasmid Construction

To generate the inducible KRAS^G12V^ expression vector, a myc-tagged KRAS^G12V^4b sequence was cloned by Gateway recombination into the tetracycline-inducible lentiviral vector pSLIK-TRE-IRES-Venus, generating pSLIK-TRE-myc-KRAS^G12V^-IRES-Venus. This construct provides bicistronic expression of KRAS^G12V^ and the fluorescent marker Venus (YFP), enabling doxycycline-inducible expression and drug-free identification of infected cells. The pHAGE-Rpl22-3X-HA-tdTomato lentiviral vector used for TRAP-seq was generated as previously described [18].

### Viral Infection and Generation of Stable Cell Lines

Lentiviruses were produced by co-transfection of the expression construct with the packaging plasmids pCMV-VCV-G and pCMV-delta into FS293 cells using the Lipofectamine 3000 Transfection Kit (Invitrogen, L3000-001). After 24 hours, medium was replaced with MCDB131 containing 20% heat-inactivated FBS. High-titer viral supernatant was harvested at 24 and 48 hours following medium change. Low-passage, sub-confluent HUVECs were infected for 18-24 hours, after which viral medium was removed and replaced with ECGM2 for at least 24 hours. Infection efficiency was assessed by Venus or tdTomato fluorescence. For double-infection experiments, HUVECs stably expressing pHAGE-Rpl22-3X-HA-tdTomato were re-seeded and infected with pSLIK-TRE-myc-KRAS^G12V^-IRES-Venus by the same protocol. For proliferation tracking in co-culture, KRAS^G12V^ cells were additionally infected with the S-G2-M phase reporter pRetroX-SG2M-Cyan (Takara Bio, #631462).

### 3D Planar Co-culture Morphogenesis Assay

The 3D planar co-culture morphogenesis assay was performed as previously described [17] with modifications as reported [12, 18]. NHDFs were seeded onto gelatin-coated 60 mm dishes 18 hours prior to endothelial cell seeding at a 1:10 ratio (HUVECs:NHDFs). Co-cultures were maintained in ECGM2 with medium changes every 2–3 days and analyzed at the indicated time points up to day 12. KRAS^G12V^ expression was induced by addition of 250 ng/mL doxycycline at the time of HUVEC seeding and maintained throughout. For TRAP-seq experiments, co-cultures were established as described and harvested on day 5 or day 12.

### Immunofluorescence and Image Analysis

For morphology assays, co-cultures were fixed at the indicated timepoints in 3.7% paraformaldehyde and stained with a rhodamine-conjugated *Ulex europaeus* agglutinin I (UEA-1) lectin to label the HUVEC cytoplasm (Vector Laboratories, FL-1061-2, 1:400). For proliferation assays in co-culture and for 2D cell size measurements, fixed and permeabilized cells (0.2% Triton X-100, 10–20 minutes) were stained with anti-ERG antibody (#16606, Cell Signaling Technology) to identify HUVEC nuclei, followed by an appropriate fluorescent secondary antibody. For 2D cell size measurements, HUVECs were induced with 50 ng/mL doxycycline for 24 hours in either serum-free medium or complete growth medium prior to fixation and staining. Total fluorescent EC area per field was quantified in ImageJ. Average area per cell was calculated by dividing total lectin-positive area by the ERG-positive cell count in the same field. Feret’s diameter was determined from lectin-stained images in ImageJ. Vessel-like structure area at day 12 was quantified using AngioTool image analysis software [19].

### Proliferation Assays

To track the proliferative index across the co-culture time course, HUVECs co-expressing KRASG12V and the S-G2-M-Cyan reporter were co-cultured with NHDFs and fixed at daily intervals in 3.7% paraformaldehyde, then permeabilized and stained for ERG as described above. The percentage of proliferating cells was calculated as the ratio of SG2M-positive to total ERG-positive nuclei per field. For 2D contact inhibition experiments, double-infected cells were seeded at sub-confluence in ECGM2 with 250 ng/mL doxycycline and analyzed at daily intervals over five days as cultures reached confluence, using the same SG2M/ERG protocol.

For EdU incorporation assays in 2D culture, HUVECs were seeded subconfluently and allowed to attach for approximately 6 hours in complete growth medium. Medium was then changed to serum-free MCDB131 and, 18 hours later, replaced with serum-free MCDB131 containing 50 ng/mL doxycycline for 24 hours. Cells were subsequently incubated with EdU for 4 hours using the Click-iT™ Plus EdU Cell Proliferation Kit for Imaging, Alexa Fluor™ 594 (Invitrogen, C10639), followed by fixation and DAPI counterstain. The ratio of EdU-positive to DAPI-positive nuclei was calculated per field. For inhibitor validation of proliferation (**Supplemental Figure 7**), the same 2D EdU protocol was used with the addition of Alpelisib (S2814, 1 µM), Trametinib (S2673, 10 nM), or Pazopanib (S3012, 1 µM) alongside doxycycline during the 24-hour induction period prior to EdU labeling. All inhibitors were purchased from Selleckchem (Houston, TX).

### Transwell Migration Assay

For assessment of 2D random migration, HUVECs were seeded onto tissue culture dishes in complete growth medium and allowed to reach confluence overnight. Medium was then replaced with serum-free medium containing 50 ng/mL doxycycline to synchronize and induce expression overnight. Transwell inserts (6.5 mm diameter, 8 µm pore polycarbonate membrane; Costar) were coated on both surfaces with 0.2% gelatin for 2 hours at 37°C, then blocked with 1% BSA in PBS for 1 hour at 37°C. Induced HUVECs were trypsinized, resuspended at 5×10⁴ cells per 100 µL in serum-free medium, and seeded into the upper chamber; serum-free medium without chemoattractant was placed in the lower chamber. Cells were allowed to migrate for 4 hours at 37°C with 5% CO₂, with rotation every 15 minutes for the first 2 hours to ensure even distribution. Inserts were fixed in 3.7% paraformaldehyde, non-migrated cells were removed from the upper surface, and migrated cells on the underside were stained with DAPI and imaged at 10× magnification. Five random fields per insert were quantified.

### Live-Cell Migration Tracking

Migration in 3D planar co-culture was assessed on day 5 by time-lapse microscopy as previously described [18]. HUVECs co-expressing the tdTomato reporter were imaged for 6 hours at 5-minute intervals using a 20× air objective on a fully automated inverted Nikon microscope equipped with Z-drift compensation, a motorized XYZ stage, a solid-state light source (Lumencor), a sCMOS camera (PCO), a heated stage, and vibration isolation, controlled through Nikon NIS-Elements software. Five fields of view were acquired per experimental condition. Individual cell tracks were generated using the mTrackJ macro plugin in Fiji/ImageJ [20, 21] and plot-at-origin analyses were generated from these tracks. Mean square displacement (MSD) and average cell speed were calculated from the same track data using the DiPer macro [22] in Excel.

### Western Blotting

HUVECs were induced with the indicated concentrations of doxycycline in serum-free medium for 24 hours prior to lysis. Cells were lysed directly in 2X Laemmli sample buffer (Bio-Rad, #1610737) supplemented with PhosSTOP EASYpack phosphatase inhibitors (Roche, #04906845001) and cOmplete Mini protease inhibitor cocktail (Roche, #53945000), then boiled at 95°C for 5 minutes. Lysates were resolved by SDS-PAGE (100 V, 1.5 hours in Tris-glycine-SDS running buffer) and transferred to 0.45 µm or 0.20 µm nitrocellulose membrane (300 mA, 1 hour in Tris-glycine/10% methanol transfer buffer). Membranes were blocked in 5% BSA with 0.02% NaN₃ for 1 hour at room temperature, then incubated overnight at 4°C with primary antibodies. After three washes in 0.05% Tween-20/TBS, membranes were incubated with goat anti-mouse IRDye 800CW and goat anti-rabbit IRDye 680RD secondary antibodies (LI-COR) for 1 hour at room temperature, washed, and imaged on the LI-COR Odyssey imaging system.

Primary antibodies and dilutions used were as follows: Myc (9E10; sc-40, Santa Cruz, 1:1000), Pan-RAS (OP40, Calbiochem, 1:1000), pAKT (Ser473; 4060S, Cell Signaling, 1:1000), AKT (pan; 2920S, Cell Signaling, 1:1000), pERK1/2 (sc-136521, Santa Cruz, 1:1000), ERK2 (sc-1647, Santa Cruz, 1:1000), pS6 (Ser235/236; 4857, Cell Signaling, 1:1000), S6 (2317S, Cell Signaling, 1:1000), GAPDH (2118S, Cell Signaling, 1:1000), cleaved caspase-3 (9661S, Cell Signaling, 1:1000), and total caspase-3 (9662, Cell Signaling, 1:1000). Phospho-protein levels were normalized to their respective total protein. For inhibitor target engagement experiments (**Supplemental Figure 6**), KRAS^G12V^ HUVECs were treated with Alpelisib (1 µM) or Trametinib (10 nM) for 24 hours in complete growth medium prior to lysis.

### TRAP-seq

#### RNA Isolation

The TRAP-seq protocol was performed as previously described [18, 23]. At the indicated harvest timepoint (day 5 or day 12), co-cultures were washed with ice-cold PBS containing 100 µg/mL cycloheximide and flash frozen. Cells were lysed in 400 µL polysome lysis buffer per 60 mm dish, homogenized, and incubated with 1% NP-40 for 20 minutes at 4°C, followed by centrifugation at 12,000 rpm for 10 minutes at 4°C. The RNA-containing supernatant was incubated with Pierce anti-HA magnetic beads (Thermo Fisher, #88837) for 5–18 hours at 4°C to immunoprecipitate HA-Rpl22-associated polysomes. RNA was eluted from beads using the Qiagen RNeasy Plus Micro Kit (#74034). RNA concentration was determined with the Qubit RNA High Sensitivity Kit (Invitrogen, #Q32853C), and RNA integrity was assessed using the Agilent RNA Pico Chip on the Agilent 2100 Bioanalyzer (#5057-1513). Only samples with RIN ≥ 7 were used for library preparation.

#### Library Preparation, Sequencing, and Alignment

RNA-seq libraries were prepared by Azenta Life Sciences and sequenced on an Illumina NextSeq 500, generating paired-end reads. Raw FASTQ files were trimmed with Trim Galore to remove adapter sequences and low-quality bases. Reads were aligned to the *Homo sapiens* reference genome (GRCh38/hg38) using the *align* function from the Rsubread package [24], and mapping efficiency was assessed with the *propmapped* function. Gene-level counts were generated with *featureCounts* in Rsubread using the inbuilt hg38 gene annotation, retaining only uniquely mapped, properly paired reads. Genes with fewer than 0.5 counts-per-million in at least 6 of the 16 libraries were excluded from further analysis; 14,716 genes passed this filter and constituted the detected gene set for all downstream analyses.

#### Differential Expression Analysis

Differential expression analysis was performed in R using DESeq2 [25]. Raw counts were modeled with the design formula ∼ vendor + condition_timepoint, where vendor (PromoCell or Lonza) was included as a covariate to account for inter-lot variation among the four independent HUVEC donor pools, and condition_timepoint encoded the four experimental groups (WT_D5, KRAS^G12V^_D5, WT_D12, KRAS^G12V^_D12). Size factors were estimated by the median-of-ratios method, and dispersions were estimated using the default DESeq2 empirical Bayes procedure.

Pairwise contrasts were extracted for KRAS^G12V^ vs. WT at day 5, KRAS^G12V^ vs. WT at day 12, and an interaction term capturing the differential change in KRAS^G12V^ effect from day 5 to day 12. P-values were adjusted by the Benjamini-Hochberg procedure. Genes with FDR < 0.05 and |log₂FC| > 0.585 were considered significant, corresponding to a minimum 1.5-fold change. Variance-stabilizing transformation (VST) was applied to normalized counts for visualization.

#### Principal Component Analysis

PCA was performed on log₂-transformed DESeq2 normalized counts (log₂[normalized counts + 0.5]) using the 500 most variable genes across all 16 samples. PC1 (45.8% of variance) separated samples by HUVEC donor lot of origin, confirming vendor as the primary source of technical variation in the dataset; PC2 (21.3% of variance) separated samples by genotype and timepoint, reflecting the biological signal under study.

#### Overrepresentation Analysis

Overrepresentation analysis (ORA) was performed by one-tailed hypergeometric test using the 14,716 detected genes as the background. Gene sets tested included the MSigDB Hallmark collection [26] and three curated sets: SASP/Secretory Phenotype (Reactome R-HSA-2559582; Coppé et al. 2008), Oncogene-Induced Senescence (Reactome R-HSA-2559583), and Hippo/YAP pathway ([27]; MSigDB C6 CORDENONSI_YAP_CONSERVED_SIGNATURE). ORA was run separately for upregulated and downregulated DEGs at each contrast, and Benjamini-Hochberg correction was applied independently within each direction. Terms with FDR < 0.05 were considered significant.

#### Curated Gene Panels

To interrogate specific biological programs not fully captured by Hallmark gene sets, three curated gene panels were assembled from the literature and used for focused analysis of the TRAP-seq data.

A migration panel of 275 genes was organized into six functional categories: Cytoskeleton/Actin Dynamics, Rho GTPase Signaling, Focal Adhesion and ECM, Growth Factor/Chemotaxis, Membrane Protrusions and Polarity, and KRAS-Direct Migration. Category assignments were based on documented gene function in EC migration and motility contexts [28-32]. Genes were included only if detected in the DESeq2 analysis. ORA within the migration panel used the same hypergeometric test and directional BH correction as described above, applied at the day 5 contrast.

A Notch pathway panel of 97 genes was organized into eight categories: Receptors, Ligands (DSL), γ-Secretase complex, Canonical Effectors, Notch Coregulators, Notch Endothelial Targets, AVM/Arteriovenous Identity, and Notch Cross-talk. Category assignments were based on established pathway annotations and the intersection of Notch signaling with EC identity and AVM biology [33, 34]. Of the 97 genes in the panel, 49 were significant at one or more contrasts and are displayed in **Supplemental Figure 5.**

A subset of 12 genes from the AVM/Arteriovenous Identity category of the Notch panel was used for focused display of AVM-related transcriptional changes across timepoints (**Figure 7B**). This set included representatives of four functional groups: TGF-β/BMP signaling, Notch/arterial identity, ECM and guidance, and VEGF/angiogenic signaling. All 12 genes reached significance at day 12.

### Statistical Analysis

All statistical analyses were performed in GraphPad Prism or in R. Experiments were conducted with at least four independent pools of primary HUVECs from two commercial vendors (PromoCell and Lonza), each pool representing cells from at least three donors of mixed sexes and ethnicities; each *n* represents one independently seeded experiment. Comparisons between two groups were made by two-tailed unpaired Student’s *t*-test. Multi-group comparisons were made by one-way ANOVA with Tukey’s post-hoc test, or by one-way ANOVA with Dunnett’s post-hoc test where all conditions were compared against a single reference group. Time-course data were analyzed by two-way ANOVA. For all sequencing analyses, p-values were adjusted by the Benjamini-Hochberg method as described above. The statistical test and sample size for each experiment are noted in the figure legends. Data are presented as mean ± SEM.

### Data Availability

TRAP-seq data are deposited in GEO (accession GSE336689). R analysis code is available from the corresponding author upon request.

## RESULTS

### Ras-related signaling and cellular functions are augmented with KRAS^G12V^ expression in 2D culture

To express the constitutively active, KRAS^G12V^, in primary human umbilical vein endothelial cells (HUVECs), we generated a lentiviral vector to stably deliver KRAS^G12V^ under the control of a doxycycline-inducible promoter. Western blot analysis of 2D-cultured HUVECs in the absence of growth factors confirmed KRAS^G12V^ protein expression approximately 3.5-fold above endogenous KRAS levels 24 hours after doxycycline induction (**Supplemental Figure 1A and 1B**). Downstream of KRAS^G12V^, phosphorylation of AKT, S6, and ERK were all elevated relative to empty vector control cells, confirming activation of both PI3K/AKT and MAPK/ERK pathways (**Supplemental Figure 1C and 1D**).

To determine how this abnormal signaling affects EC function in 2D culture, we examined proliferation, migration, and survival. In serum-free conditions, KRAS^G12V^ cells exhibited significantly higher EdU incorporation compared to control, though this difference was lost in the presence of a full complement of growth factors (**Supplemental Figure 2A**). KRAS^G12V^ also enhanced random migration across a transwell membrane under serum-free conditions (**Supplemental Figure 2B**). To assess survival, we measured cleaved caspase-3 (CC3) after 72 hours in serum-free medium and found significantly lower CC3 levels in KRAS^G12V^ cells compared to control, demonstrating a survival advantage in the absence of growth factors (**Supplemental Figure 2C**). Consistent with *in vivo* reports of KRAS^G12V^-driven EC hypertrophy, we also found that KRAS^G12V^ cells had increased Feret’s diameter and average area per cell in both serum-free and complete growth conditions (**Supplemental Figure 3A–C**). Together, these observations in 2D culture establish that KRAS^G12V^ is sufficient to alter several EC behaviors that could contribute to abnormal vascular morphogenesis.

### KRAS^G12V^ disrupts normal morphogenesis and drives a transient proliferative phase in 3D angiogenesis assays

To examine the morphological consequences of KRAS^G12V^ expression in a context more closely resembling vascular development, we seeded HUVECs in the 3D planar co-culture assay with primary normal human dermal fibroblasts (NHDFs) and analyzed cultures by fluorescent lectin staining at daily intervals from day 1 through day 12 (**Figure 1A**). Control cells progressively organized into thin, branched, vessel-like networks. KRAS^G12V^ cells instead formed a compact, sheet-like morphology that diverged from control structures as early as day 3 and became increasingly abnormal through day 12. Quantification of total fluorescent area confirmed a statistically significant divergence between genotypes beginning on day 3 and persisting across all subsequent time points (**Figure 1B**). At day 12, control co-cultures contained abundant tubular structures detectable by AngioTool analysis, whereas KRAS^G12V^ co-cultures showed a near-complete absence of vessel-like structures (**Figure 1C**).

**Figure 1.**
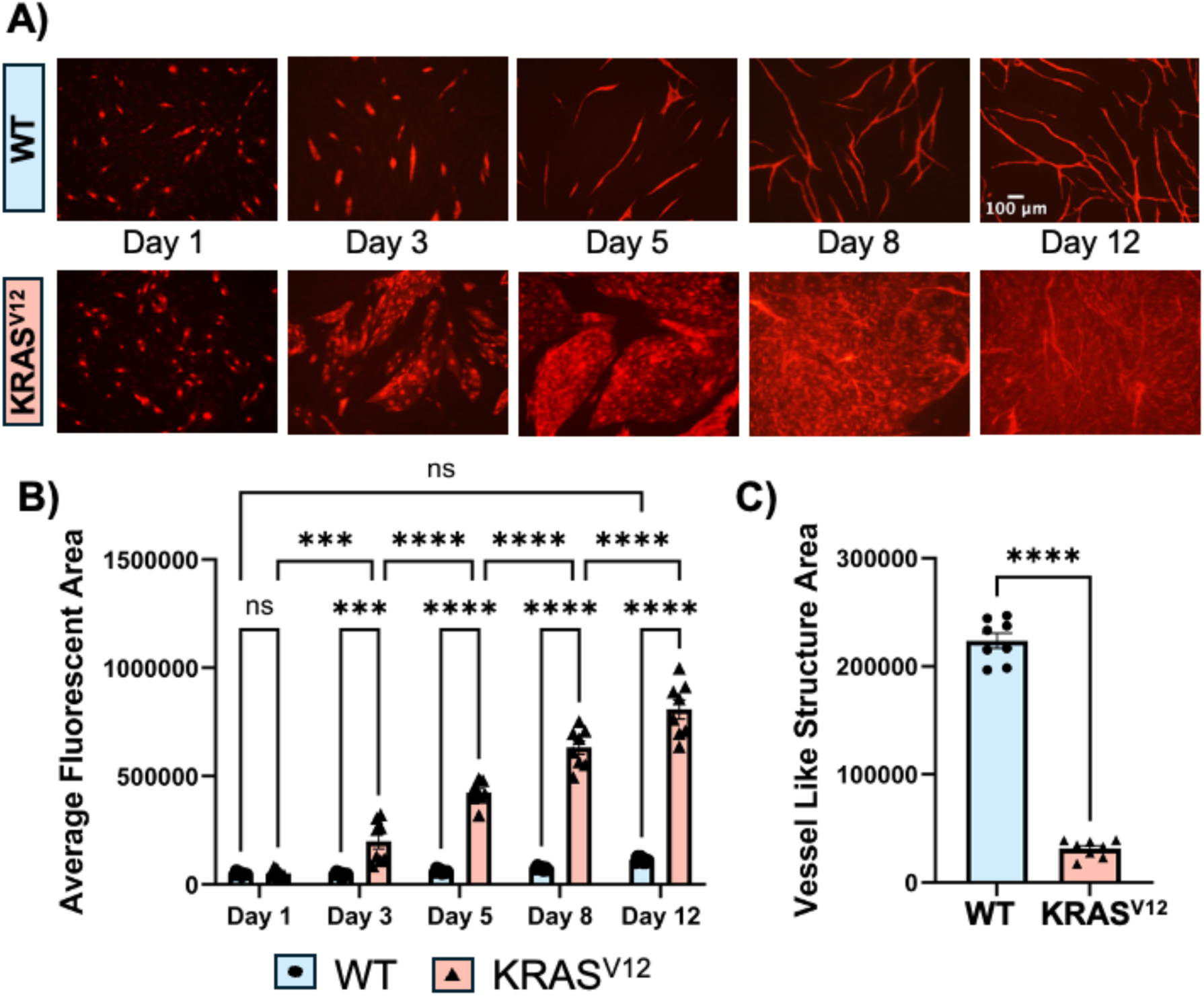
KRAS^G12V^ disrupts normal morphogenesis in 3D angiogenesis assays. A) Representative images of HUVECs expressing an empty vector control (WT) or KRAS^G12V^ from day 1 to day 12 of the planar co-culture assay with primary NHDFs at 250 ng/mL doxycycline. Cells were fixed at each timepoint and visualized with a fluorescent RFP-lectin stain. WT cells progressively organized into thin, branched vessel-like networks, while KRAS^G12V^ cells formed a compact, sheet-like morphology that diverged from controls from day 3 onward. Images are representative of 8 independent experiments. Scale bar = 100 μm; scale bar applies to all images. B) Quantification of total fluorescent area per field of view across the D1–D12 time course for the experiments represented in A. Total fluorescent area reflects both tubular structures and sheets. KRAS^G12V^ cells showed significantly greater fluorescent area beginning at day 3 and persisting through day 12. N=8 independent experiments. Two-way ANOVA, p<0.05, p<0.01, p<0.001, p<0.0001. C) Quantification of vessel-like structure area at day 12 as determined by AngioTool image analysis software. Control co-cultures contained abundant vessel-like structures at day 12, whereas KRAS^G12V^ co-cultures showed a near-complete absence of such structures. N=8 independent experiments. Student’s T-test, p<0.0001. Data represent the mean and SEM.

The marked expansion of total lectin-positive area in KRAS^G12V^ co-cultures, reflecting a larger EC sheet rather than organized tubular structures, suggested that enhanced EC proliferation might contribute to this sheet-like phenotype. To test this directly, we used a fluorescent SG2M cell-cycle reporter that is expressed during the S, G2, and M phases and degraded upon entry into G0/G1 [18, 35]. KRAS^G12V^ cells co-infected with this reporter were analyzed across the full-time course alongside ERG staining to identify the total EC population (**Figure 2A**). The proportion of proliferating KRAS^G12V^ cells was significantly elevated relative to control at day 3, remained elevated through day 5, and then declined progressively, becoming indistinguishable from control by day 12 (**Figure 2B**). Total ERG counts increased correspondingly over the proliferative phase before plateauing (**Figure 2C**). This proliferative behavior was therefore transient, occurring primarily between days 3 and 5, followed by growth arrest by day 12.

**Figure 2.**
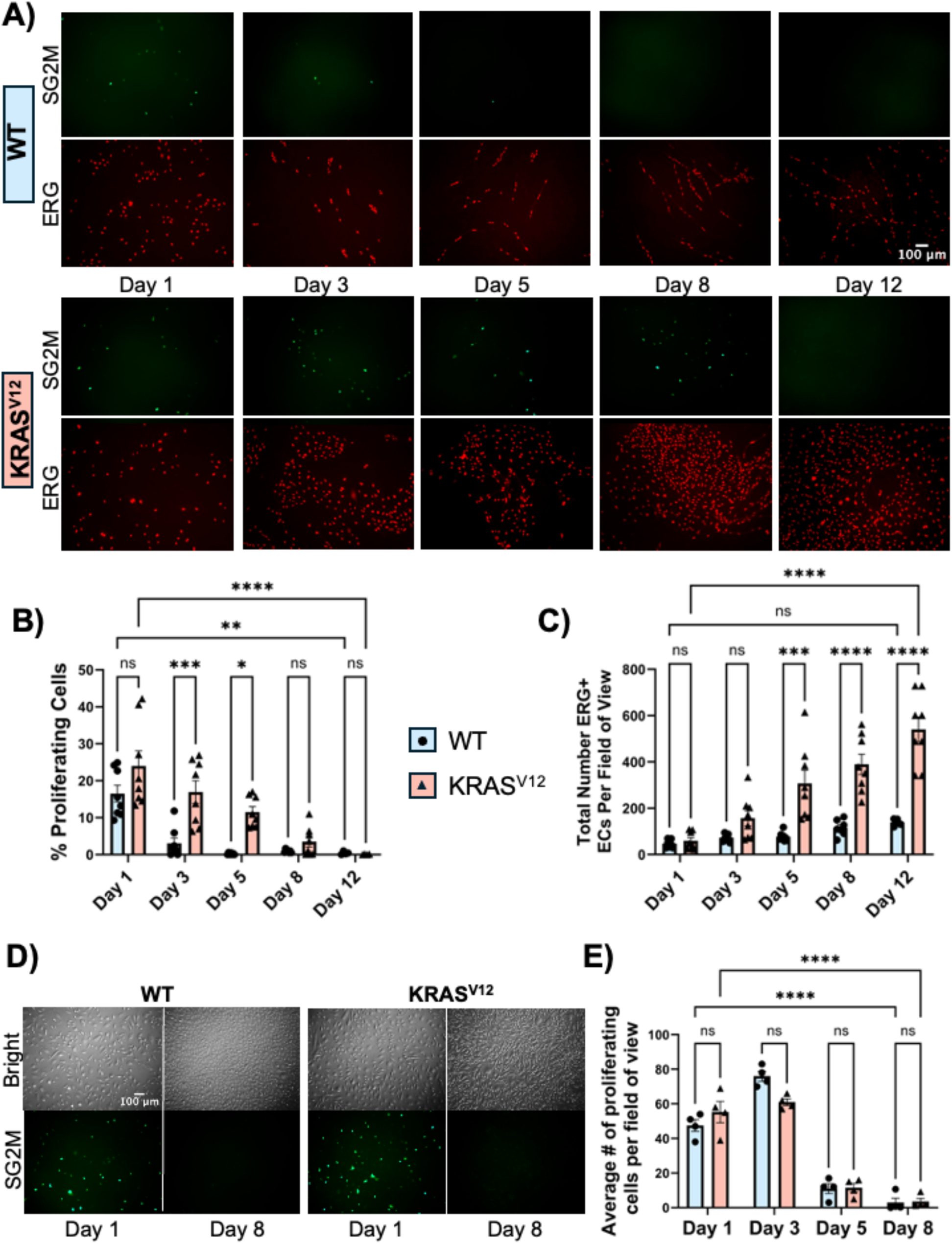
KRAS^G12V^ expression is associated with a transient proliferative phase in 3D planar co-culture; KRAS^G12V^ cells remain sensitive to contact inhibition. A) Representative images of HUVECs co-expressing WT vector (top rows) or KRAS^G12V^ (bottom rows) with the fluorescent SG2M cell cycle reporter from day 1 to day 12 of the planar co-culture assay at 250 ng/mL doxycycline. Cultures were fixed at each timepoint and stained with ERG to identify the total endothelial cell population. Images are representative of 8 independent experiments. Scale bar = 100 μm. B) Quantification of the proportion of proliferating cells (SG2M-positive / ERG-positive) at each timepoint. KRAS^G12V^ cells showed significantly elevated proliferation relative to control at day 3 and day 5. The proportion declined progressively and was indistinguishable from control by day 12. N=8 independent experiments. Two-way ANOVA; Student’s T-test at individual timepoints. C) Quantification of total ERG-positive cell count per field of view across the time course. KRAS^G12V^ cultures accumulated significantly more cells beginning at day 5 and continuing through day 12. N=8 independent experiments. Two-way ANOVA, p<0.05, p<0.01, p<0.001, p<0.0001. D) Representative brightfield and SG2M fluorescence images of WT (left) or KRAS^G12V^ (right) HUVECs co-expressing the SG2M reporter in 2D culture with a full complement of growth factors, at day 1 (sub-confluence) and day 8 (post-confluence). Additional images for quantification were taken at days 3 and 5. Images are representative of 4 independent experiments. Scale bar = 100 μm. E) Quantification of SG2M-positive cell count per field of view at days 1, 3, 5, and 8. Both WT and KRAS^G12V^ cells showed equivalent and significant declines in proliferation as cultures reached contact confluence, with near-complete growth arrest by day 8 in both genotypes. N=4 independent experiments. Two-way ANOVA, p<0.05, p<0.01, p<0.001, p<0.0001. Data represent the mean and SEM.

The cessation of proliferation at late time points raised the question of whether KRAS^G12V^ cells retain normal contact inhibition or whether another mechanism arrests growth as the sheet matures. To address this, we allowed double-infected KRAS^G12V^/SG2M reporter cells to grow to confluence in 2D culture with a full complement of growth factors (**Figure 2D**). The proportion of proliferating cells declined significantly as cultures reached confluence and was nearly abolished within 5 days of achieving contact, in a pattern indistinguishable from control (**Figure 2E**). These data indicate that KRAS^G12V^ cells remain fully responsive to contact inhibition. The transient proliferative burst observed in the 3D co-culture is therefore more consistent with a context-dependent delay in the onset of the contact inhibition signal, likely consequent to the sheet morphology itself, rather than a fundamental override of quiescence machinery.

### KRAS^G12V^ endothelial cells exhibit enhanced migration in 3D co-culture

Given that the sheet-like morphology with KRAS^G12V^ could reflect not only abnormal proliferation but also altered migratory behavior, we assessed EC migration directly using live-cell imaging. Representative plot-at-origin track overlays show that KRAS^G12V^ cells exhibited substantially longer migratory tracks than control cells, which remained largely stationary over the same period (**Figure 3A**). Quantification of mean square displacement (MSD) at the end of the 6-hour imaging period confirmed a significant increase in displacement with KRAS^G12V^ relative to control (**Figure 3B**). These results establish that KRAS^G12V^ expression is sufficient to enhance EC migration during active morphogenesis in 3D co-culture.

**Figure 3.**
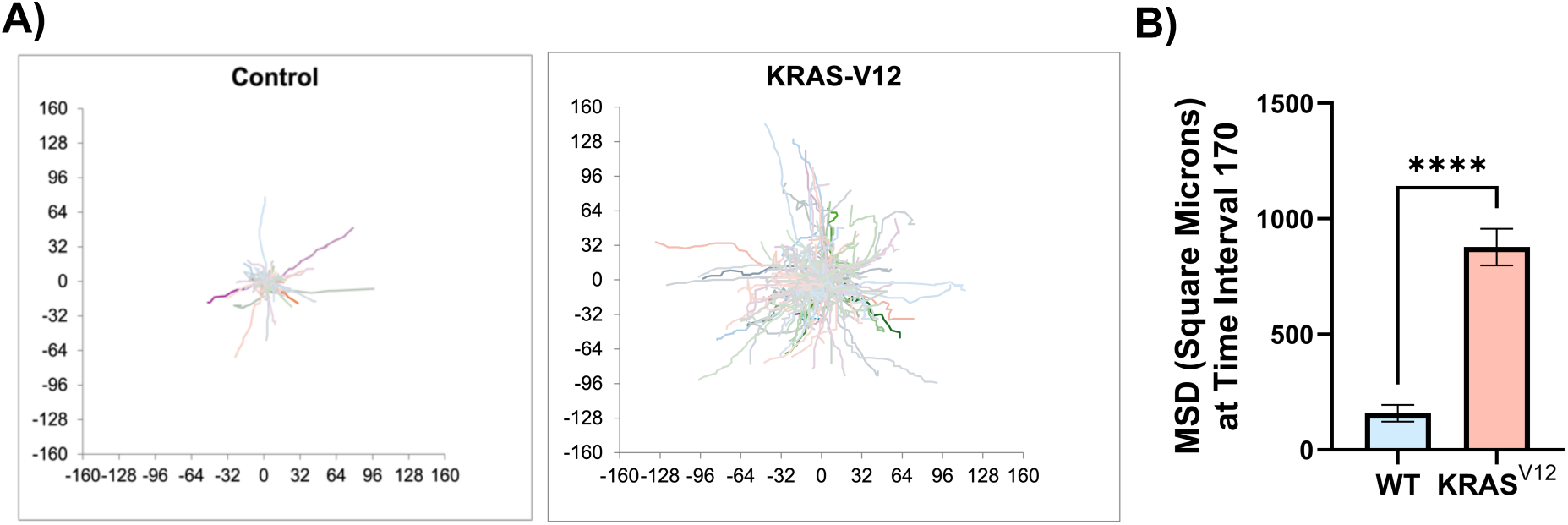
Live-cell migration tracking of endothelial cells in day 5 planar co-cultures shows enhanced migration with KRAS^G12V^. A) Representative plot-at-origin track overlays for WT and KRAS^G12V^ HUVECs at 250 ng/mL doxycycline, imaged for 6 hours at 5-minute intervals on day 5 of the planar co-culture assay. Individual cell tracks were generated using the mTrackJ macro in ImageJ. KRAS^G12V^ cells displayed substantially longer migratory tracks than WT cells, which remained largely stationary over the same period. B) Quantification of mean square displacement (MSD) in square microns at time interval 175. KRAS^G12V^ cells showed significantly greater MSD than WT controls. Representative of 4 independent seedings; tracks measured for WT N=127 cells and KRAS^G12V^ N=186 cells. Student’s T-test, p<0.0001. Data represent the mean and SEM.

### TRAP-seq reveals broad transcriptional reprogramming in KRAS^G12V^ endothelial cells across morphogenesis

To characterize the transcriptional program driving the morphological and behavioral phenotypes, we isolated HUVEC-specific mRNA from day 5 and day 12 planar co-cultures by TRAP-seq [18]. Differential gene expression analysis was performed in R using DESeq2 with a design that included HUVEC vendor as a covariate to account for inter-lot variation, with BH correction and a significance threshold of FDR < 0.05 and absolute log₂FC > 0.585.

Principal component analysis of log₂-transformed normalized counts across all 16 samples showed clean separation of samples by genotype and timepoint along PC2 (21.3% of variance), confirming that KRAS^G12V^ drives a reproducible transcriptional response across independent donor pools and vendors (**Figure 4A**). PC1 (45.8% of variance) reflected the donor lot of origin, separating PromoCell from Lonza samples. The top PC1-loading genes included proliferation-associated transcripts, consistent with a difference in proliferative state at the time of vendor cryopreservation rather than culture-induced drift, since all experiments were performed with low-passage cells purchased directly from the vendor (**Supplemental Figure 4A**). This donor-lot effect was incorporated as a covariate in the DESeq2 design, and inter-vendor expression concordance remained high at both time points (r = 0.960 at Day 5, r = 0.967 at Day 12), confirming effective correction (**Supplemental Figure 4B, C**).

**Figure 4.**
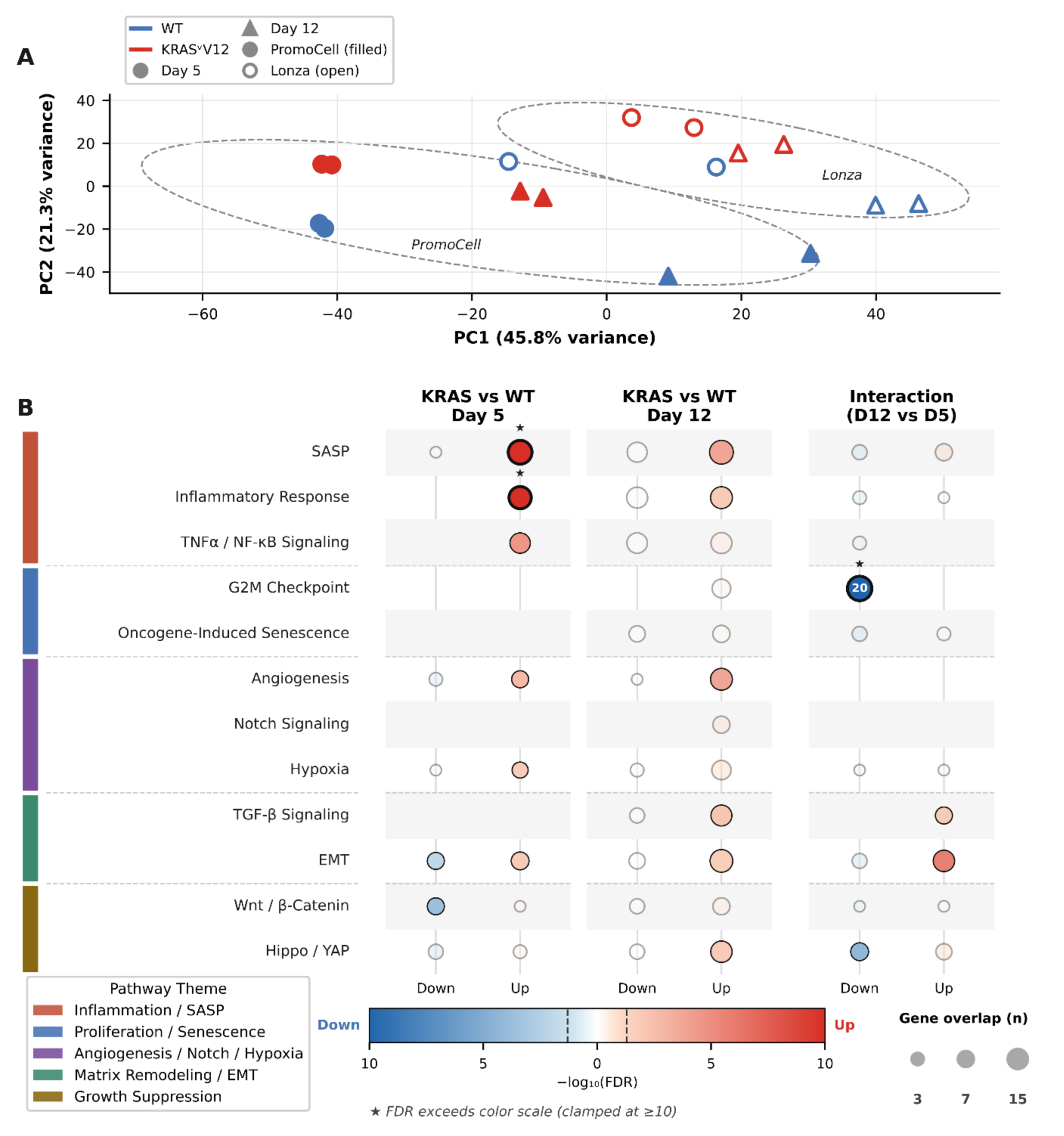
KRAS^G12V^ drives broad transcriptional reprogramming in endothelial cells revealed by TRAP-seq. A) Principal component analysis of TRAP-seq samples. Each point represents one sample; color indicates genotype (blue, WT; red, KRAS^G12V^), shape indicates timepoint (circle, Day 5; triangle, Day 12), and fill indicates HUVEC vendor lot (filled, PromoCell; open, Lonza). Dashed ellipses denote 1.8 SD confidence regions. PC1 (45.8% of variance) separates samples by vendor; PC2 (21.3%) separates by genotype and timepoint. B) Pathway enrichment landscape across three contrasts: KRAS^G12V^ vs. WT at Day 5 (320 upregulated, 202 downregulated genes), KRAS^G12V^ vs. WT at Day 12 (1,433 upregulated, 1,580 downregulated), and an interaction term capturing how the KRAS^G12V^ effect changes between Day 5 and Day 12 (338 upregulated, 151 downregulated). Each row represents a pathway grouped by biological theme (sidebar); columns represent downregulated and upregulated directions within each contrast. Bubble color encodes −log₁₀(FDR) on a diverging blue-to-red scale (blue, downregulated in KRAS^G12V^; red, upregulated); bubble size encodes gene overlap count; faded bubbles indicate pathways that did not reach significance (FDR ≥ 0.05). ★ denotes FDR values exceeding the color scale ceiling (−log₁₀(FDR) ≥ 10). N = 4 independently seeded co-cultures per genotype per timepoint.

To characterize how the KRAS^G12V^ transcriptional program evolves across morphogenesis, we performed ORA against MSigDB Hallmark and curated gene sets at three levels: KRAS^G12V^ vs. WT at day 5, KRAS^G12V^ vs. WT at day 12, and an interaction term capturing how the KRAS^G12V^ effect changes between timepoints, displayed across a unified pathway landscape (**Figure 4B**). Pathways were grouped into five biological themes: Inflammation/SASP, Proliferation/Senescence, Angiogenesis/Notch, Matrix Remodeling/EMT, and Growth Suppression (**Figure 4B**).

#### KRAS^G12V^ promotes acute inflammatory and migratory programs in day 5 co-cultures

At day 5, the most prominent upregulated gene sets were inflammatory: *SASP* (senescence-associated secretory phenotype) (Fold Enrichment (FE) = 13.4, padj = 6.9×10⁻¹⁵, n = 18 genes), *Inflammatory Response* (FE = 11.5, padj = 6.1×10⁻¹¹, n = 14), and *TNFα/NF-κB signaling* (FE = 5.8, padj = 3.5×10⁻⁵, n = 10). Hallmark *Angiogenesis* was also significantly enriched (FE = 6.8, padj = 0.002, n = 5) and examined at the individual gene level in ***Figure 6***. *EMT* (Endothelial-to-mesenchymal transition) genes were enriched in both the up and down-regulated directions, and *Wnt/β-Catenin* signaling was significantly suppressed (FE = 13.5, padj = 2.3×10⁻⁴, n = 5). *G2M Checkpoint*, *Oncogene-induced senescence* (OIS), *Notch Signaling*, and *Hippo/YAP* showed no significant hits at day 5 (**Figure 4B**).

We note that OIS did not reach significance for any contrast (**Figure 4B**), and that the SASP co-enrichment on day 5 occurs in cells that are actively proliferating (*as seen in Figure 2*). In OIS, the secretory phenotype accompanies growth arrest rather than active division; the co-occurrence of SASP and proliferation here is more consistent with an NF-κB-driven inflammatory response downstream of oncogenic Ras. Together, these two observations argue against a classical oncogene-induced senescence interpretation. inflammatory response downstream of oncogenic Ras. Together, these two observations argue against a classical oncogene-induced senescence interpretation.

The day 5 live-cell migration data prompted us to examine in detail the transcriptional basis of the enhanced migration phenotype. We assembled a curated panel of 275 endothelial cell motility genes, organized into six functional categories, and identified 30 genes from this panel that reached significance on day 5 (**Figure 5A**). ORA of these categories against the day 5 upregulated gene list identified three categories as significantly enriched: *Focal Adhesion & ECM* (FE = 6.57, FDR = 1.6×10⁻⁴, n = 8 genes*), KRAS-Direct Migration* (FE = 4.89, FDR = 0.010, n = 5 genes), and *Growth Factor/Chemotaxis* (FE = 4.72, FDR = 0.020, n = 4 genes) (**Figure 5B**). *Rho GTPase Signaling* was not enriched in either direction, indicating that canonical Rho/Rac cytoskeletal machinery is transcriptionally quiescent at day 5. This pattern suggests that the migratory phenotype is driven principally by matrix remodeling and chemotactic signaling rather than direct cytoskeletal reprogramming.

**Figure 5.**
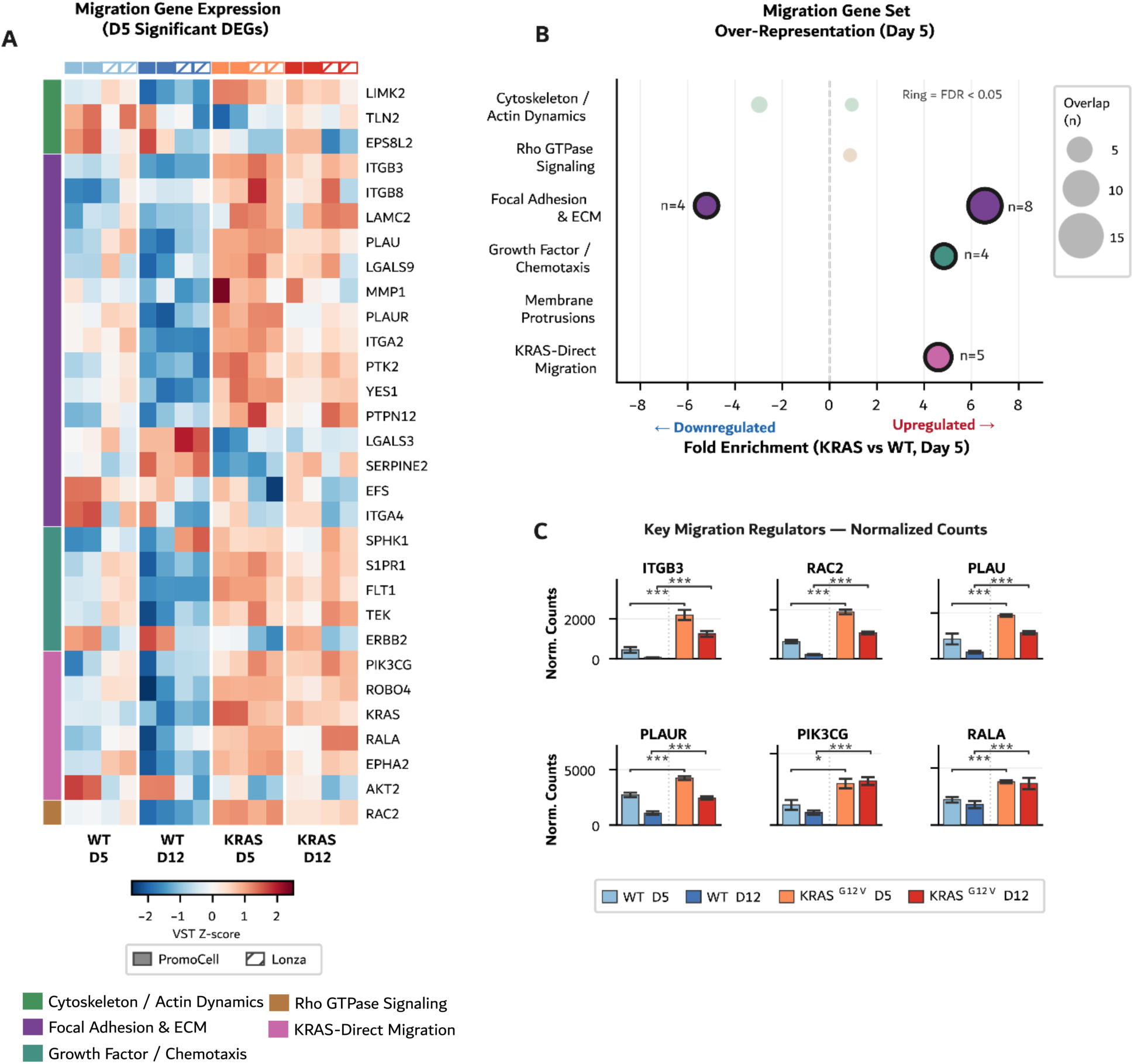
KRAS^G12V^ selectively activates matrix remodeling and chemotactic signaling pathways to drive enhanced endothelial cell migration. A) Heatmap of variance-stabilizing transformation (VST) Z-scores for the 30 migration-associated genes reaching significance at Day 5 (KRAS^G12V^ vs. WT), drawn from a curated panel of 275 endothelial cell motility genes organized into six functional categories (color sidebar). Genes within each category are ordered by descending Day 5 log₂FC. Columns represent individual samples ordered by group (WT Day 5, WT Day 12, KRAS^G12V^ Day 5, KRAS^G12V^ Day 12); solid annotation bars indicate PromoCell donor lots; hatched bars indicate Lonza donor lots. Color scale: RdBu_r, clipped at ±2.5. B) Over-representation analysis of each functional category against the Day 5 upregulated gene list. Bubble x-position indicates fold enrichment; bubble size indicates the number of overlapping genes; black rings indicate FDR < 0.05. Three categories reached significance in the upregulated direction: Focal Adhesion & ECM (FE = 6.57, FDR = 1.6×10⁻⁴, n = 8 genes), KRAS-Direct Migration (FE = 4.89, FDR = 0.010, n = 5), and Growth Factor/Chemotaxis (FE = 4.72, FDR = 0.020, n = 4). Rho GTPase Signaling was not enriched in either direction, indicating that canonical Rho/Rac cytoskeletal machinery is transcriptionally quiescent at Day 5. C) Normalized count expression of six key migration regulators, *ITGB3*, *PLAU*, *PLAUR*, *RAC2*, *PIK3CG*, and *RALA*, in WT and KRAS^G12V^ endothelial cells at Day 5 and Day 12. All six genes were significantly upregulated at both timepoints. Bars represent mean ± SEM; N = 4 independently seeded co-cultures. Significance brackets reflect DESeq2 adjusted p-values from DESeq2 pairwise contrasts (KRAS^G12V^ vs. WT at each timepoint): p < 0.05; p < 0.01; p < 0.001.

Examination of individual gene expression supported this interpretation (**Figure 5C**). The *Focal Adhesion & ECM* category included *ITGB3*, encoding the β3 integrin subunit with well-established roles in EC migration through VEGF-driven upregulation of αvβ3, Src-dependent adhesion signaling, and microtubule-dependent focal adhesion remodeling [36-38]. Both *PLAU* and *PLAUR*, which encode the urokinase plasminogen activator and its receptor, suggest coordinated activation of the uPA/plasmin proteolytic axis that degrades extracellular matrix to facilitate EC invasion [39]. Among the KRAS-Direct Migration genes, *PIK3CG*, encoding the p110γ catalytic subunit of PI3Kγ, was prominently elevated, as was *RALA*, a downstream Ras effector that promotes non-directional motility through exocyst-mediated vesicle trafficking.

*RAC2* was the sole Rho/Rac family member to reach significance at day 5, consistent with the ORA result and pointing toward an isoform-restricted rather than pathway-wide GTPase response. All six genes were significantly upregulated at both day 5 and day 12, indicating a sustained rather than transient migratory program (**Figure 5C**). The complete migration gene panel with category assignments and differential expression statistics for all contrasts is provided in **Supplemental Table 2**.

Among the remaining day 5 upregulated terms, the Hallmark *Hypoxia* gene set also reached significance (FE = 4.84, padj = 0.018, n = 4 genes: *ADORA2B*, *CXCL8*, *HMOX1*, *PLOD2*) (**Figure 4B**). Three of these four genes (*HMOX1*, *CXCL8*, and *ADORA2B*) have established roles in EC survival, motility, and inflammatory signaling under vascular stress [40-42] and *PLOD2* is a HIF-1 target that modifies collagen crosslinking to alter extracellular matrix stiffness. The modest size of the hit, the overlap with inflammatory gene sets already reaching significance, and the absence of *Hypoxia* enrichment at Day 12 suggest this reflects a general cellular stress or inflammatory response at the early time point rather than a sustained or canonical hypoxic transcriptional program.

### Notch pathway genes are over-represented among Day 5 angiogenesis-associated DEGs

Among the upregulated terms in the day 5 ORA, the Hallmark Angiogenesis gene set reached significance with 5 genes (FE = 6.8, padj = 0.002). To understand which specific angiogenesis-associated genes were dysregulated, we examined each of the 5 significantly upregulated members of this gene set: *JAG1*, *ROBO4*, *FLT1*, *SPHK1*, and *MDK* (**Figure 6**). Of these 5 genes, 3 (*JAG1*, *ROBO4*, and *FLT1*) are components of the Notch signaling pathway, while the remaining 2 (*SPHK1* and *MDK*) are non-Notch angiogenesis regulators. The representation of Notch pathway components in this relatively small hit set prompted a broader, more systematic examination of the Notch transcriptional program at both time points.

**Figure 6.**
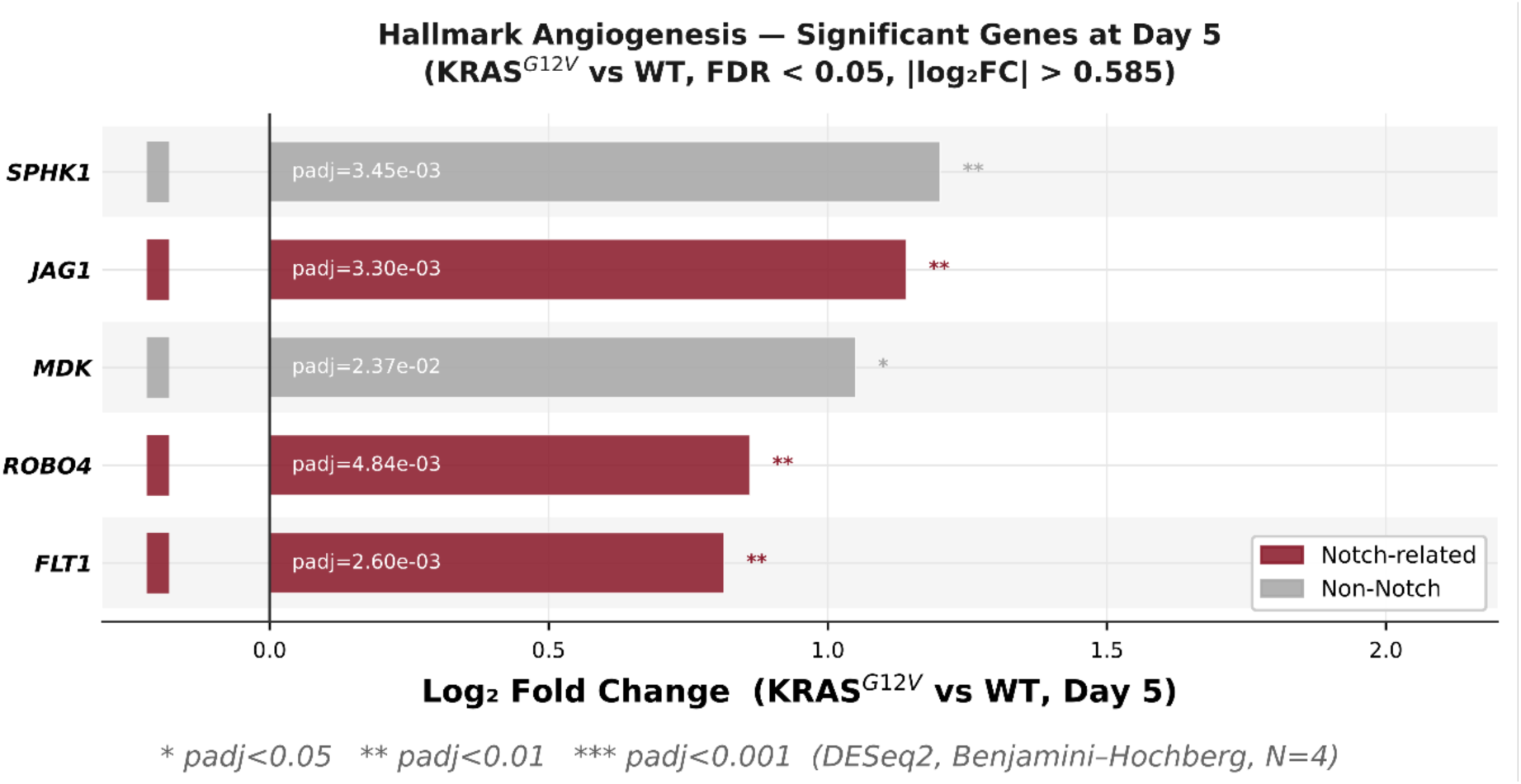
KRAS^G12V^ upregulates Notch pathway genes within the Hallmark Angiogenesis gene set at Day 5. Bar plot showing the five genes from the MSigDB Hallmark Angiogenesis gene set that are significantly differentially expressed in KRAS^G12V^ vs. WT endothelial cells at Day 5 (FDR < 0.05, DESeq2). Three of the five genes, *JAG1* (log₂FC = 1.14, padj = 0.003), *ROBO4* (log₂FC = 0.86, padj = 0.005), and *FLT1* (log₂FC = 0.81, padj = 0.003), are components of the Notch signaling pathway (maroon). The remaining two genes, *SPHK1* (log₂FC = 1.20, padj = 0.003) and *MDK* (log₂FC = 1.05, padj = 0.024), are non-Notch angiogenesis regulators (gray). The representation of Notch pathway components within this small hit set prompted a broader examination of the Notch transcriptional program at both timepoints (Figure 7). Bars represent log₂ fold change (KRAS^G12V^ vs. WT). RNA was isolated from N = 4 independently seeded Day 5 co-cultures by TRAP-seq. padj < 0.05, padj < 0.01, padj < 0.001 (Benjamini–Hochberg correction).

### KRAS^G12V^ endothelial cells acquire an AVM transcriptional identity as wild-type cells quiesce

By day 12, control ECs have completed morphogenesis and entered quiescence. The acute inflammatory signature that dominated day 5 resolved substantially: TNFα/NF-κB and Wnt/β-Catenin, both prominent at day 5, were no longer significant. In their place, six gene sets were enriched in the upregulated direction: Hallmark Angiogenesis (FE = 3.9, padj = 1.9×10⁻⁴, n = 13 genes; expanded from 5 genes at day 5), SASP (FE = 3.0, padj = 1.9×10⁻⁴, n = 18), TGF-β signaling (FE = 2.8, padj = 0.009, n = 11), Hippo/YAP (FE = 2.4, padj = 0.015, n = 12), Inflammatory Response (FE = 2.4, padj = 0.014, n = 13), and EMT (FE = 2.0, padj = 0.024, n = 15) (**Figure 7A**). The persistence and expansion of the angiogenesis signature at a point when WT cells have exited this program suggests that KRAS^G12V^ ECs fail to transition normally into quiescence.

**Figure 7.**
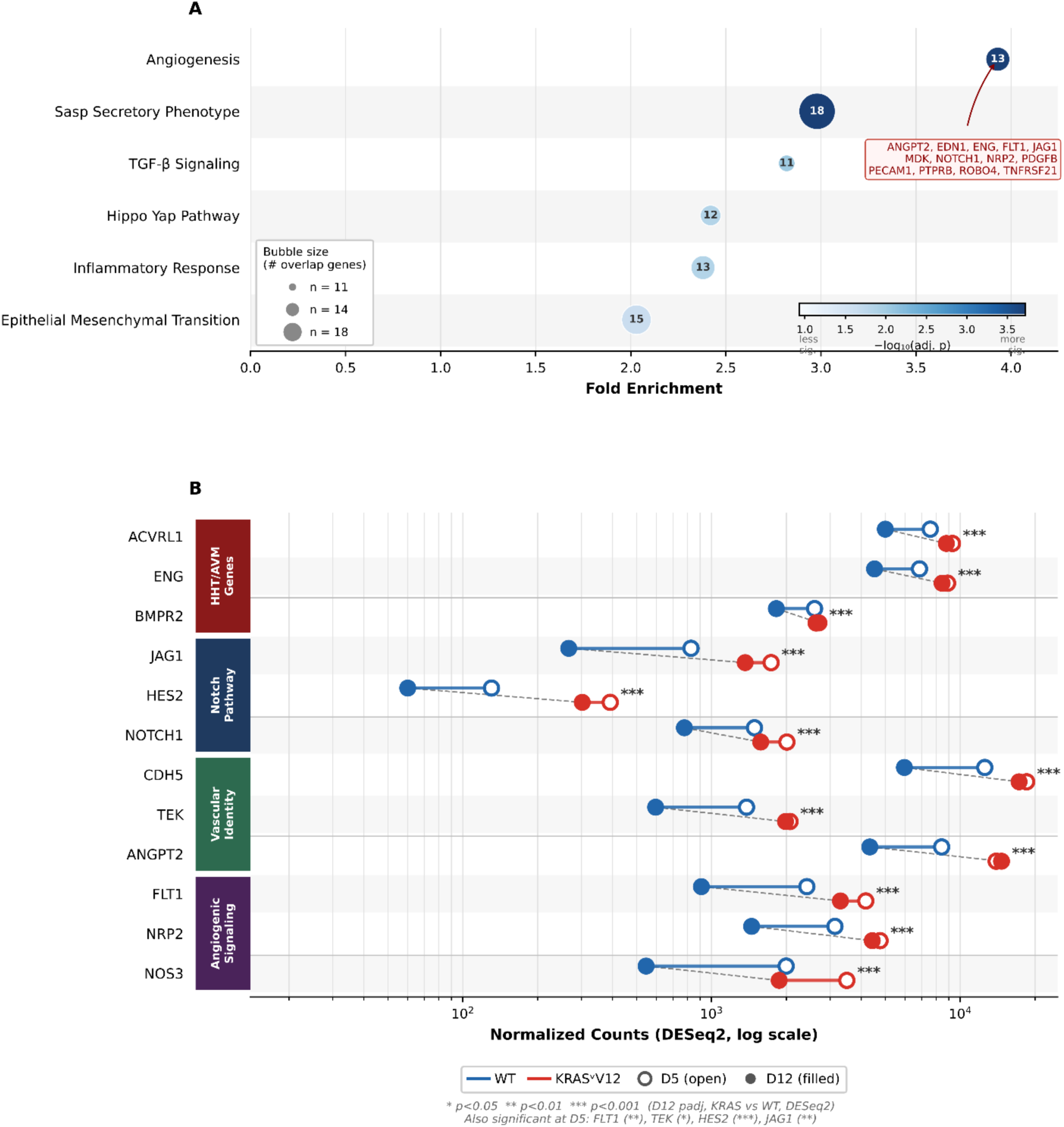
KRAS^G12V^ endothelial cells maintain an AVM transcriptional identity as wild-type cells quiesce. A) Over-representation analysis (ORA) of DESeq2-significant genes upregulated in KRAS^G12V^ vs. WT endothelial cells at Day 12 (N = 4 independent HUVEC pools). Each bubble represents one gene set; bubble size reflects overlapping gene count; bubble color encodes −log₁₀(adjusted p-value). Six terms reached significance in the upregulated direction. Hallmark Angiogenesis showed the strongest enrichment (FE = 3.93, padj = 1.9×10⁻⁴, n = 13 genes). The overlapping genes are annotated on the figure; six (ENG, JAG1, NOTCH1, ANGPT2, FLT1, NRP2) carry established roles in HHT and AVM pathobiology and are examined further in Panel B alongside six additional AVM-associated genes significantly upregulated at Day 12 (ACVRL1, BMPR2, HES2, CDH5, TEK, NOS3). Enrichment statistics for all significant terms are provided in the Results. B) DESeq2 size-factor normalized expression of AVM identity genes in WT (blue) and KRAS^G12V^ (red) endothelial cells at Day 5 (open circles) and Day 12 (filled circles) in fibroblast co-culture. Each horizontal track represents the mean of N = 4 independent HUVEC pools. Genes are grouped by functional category (color sidebar): HHT/AVM genes (*ACVRL1*, *ENG*, *BMPR2*), Notch pathway (*JAG1*, *HES2*, *NOTCH1*), vascular identity (*CDH5*, *TEK*, *ANGPT2*), and angiogenic signaling (*FLT1*, *NRP2*, *NOS3*). Dashed vertical connectors link WT and KRAS^G12V^ values at Day 12. Significance symbols indicate Day 12 adjusted p-values (KRAS^G12V^ vs. WT, DESeq2): p < 0.05, p < 0.01, p < 0.001, p < 0.0001. Genes also reaching significance at Day 5: HES2 (), *JAG1*, *FLT1*, *TEK*. Note: *BMPR2* log₂FC = 0.574 falls marginally below the |log₂FC| > 0.585 threshold applied throughout; it is retained on the basis of its padj = 3.6×10⁻⁴ and its established role in HHT/AVM pathobiology.

Several of the 13 genes underlying the Hallmark Angiogenesis enrichment are components of the Notch signaling pathway or carry established roles in HHT and AVM pathobiology, including *ENG*, *JAG1*, *NOTCH1*, *ANGPT2*, *FLT1*, and *NRP2*. To assess whether this represented a broader shift in Notch-dependent vascular identity, we examined expression across a curated panel of 97 Notch-associated genes organized into eight functional categories (**Supplemental Figure 5**). Of 47 genes reaching significance in at least one contrast, enrichment was concentrated in the AVM/arteriovenous identity and endothelial target categories, with the canonical receptor-ligand processing machinery largely unaffected. This pattern is consistent with selective activation of the vascular identity output of Notch signaling rather than pathway-wide induction and motivated a more focused examination of genes with direct AVM relevance. The complete Notch pathway panel with category assignments and differential expression statistics for all three contrasts is provided in Supplemental Table 3.

We therefore tracked expression from day 5 to day 12 across a curated panel of 12 genes in four functional categories drawn from the HHT and AVM literature (**Figure 7B**). Six of these genes (*ENG*, *JAG1*, *NOTCH1*, *ANGPT2*, *FLT1*, *NRP2*) are members of the Hallmark Angiogenesis set; the remaining six (*ACVRL1*, *BMPR2*, *HES2*, *CDH5*, *TEK*, *NOS3*) are significantly upregulated at day 12 but fall outside the Hallmark gene set definition, and were included on the basis of their established roles in HHT/AVM pathobiology [43]. The majority of these genes were not significantly changed at day 5 but became strongly elevated by day 12 as WT expression declined. *ACVRL1* (ALK1), *ENG* (endoglin), and *BMPR2*, which encode components of the TGF-β/BMP signaling axis with well-characterized roles in HHT and vascular AVM [44, 45], reached significance only at day 12 (all padj < 0.001). Early Notch activation was apparent in *JAG1* and *HES2*, both significantly elevated at day 5 and increased further by day 12; *NOTCH1* followed a similar trajectory but reached significance only at the later timepoint. *TEK* and *FLT1* were significantly elevated at both time points, whereas CDH5, ANGPT2, NRP2, and NOS3 were significant only at day 12. The upregulation of arterial identity markers *NOTCH1*, *TEK*, and *NRP2* in venous-derived HUVECs is consistent with progressive acquisition of an arterial/AVM transcriptional state under sustained KRAS^G12V^ expression.

To assess whether the day 12 KRAS^G12V^ transcriptional profile bears quantitative resemblance to human bAVM endothelium, we compared our findings directly against the nidus endothelial signature from the Winkler 2022 single-cell atlas of surgically resected human bAVM tissue [46]. Of the 632 Winkler Nd2 upregulated DEGs detected in the present dataset, 122 are significantly upregulated in KRAS^G12V^ ECs at Day 12 (fold enrichment = 1.98, p = 5.85×10⁻¹⁴, hypergeometric test; **Supplemental Table 4**), with MFSD2A and TBX2 simultaneously downregulated. Applying the GSEA approach used by Winkler et al. to our Day 12 ranked gene list, the same four Hallmark pathways they reported in Nd2 are independently enriched: Angiogenesis (NES = 1.73, padj = 0.024), Inflammatory Response (NES = 2.02, padj = 2.3×10⁻⁶), EMT (NES = 1.50, padj = 7.3×10⁻³), and TGF-β Signaling (NES = 2.05, padj = 1.7×10⁻⁴). Together, these analyses establish a quantitative resemblance between the day 12 KRAS^G12V^ transcriptional profile and human bAVM nidus endothelium at both the gene and pathway level (**Supplemental Table 4**).

#### Delayed exit from the proliferative program in KRAS^G12V^ endothelial cells

The most informative contrast was the interaction term, which captures transcriptional changes in KRAS^G12V^ relative to WT that are specifically amplified or extinguished between day 5 and day 12. The dominant signal was downregulation of the G2M Checkpoint gene set (FE = 35.4, padj = 2.4×10⁻²⁵, n = 20 genes). Examination of individual G2M genes revealed that these transcripts are expressed at comparable levels in KRAS^G12V^ and WT cells on day 5 and are then selectively retained in KRAS^G12V^ as WT cells downregulate them between day 5 and day 12 (**Supplemental Table 1, Section B**). This pattern is consistent with the phenotypic arrest of proliferation by day 12 and supports the interpretation of delayed morphogenetic quiescence: WT cells exit the proliferative transcriptional program on schedule, while KRAS^G12V^ cells do so with a lag. The Hippo/YAP pathway was also significantly downregulated in the interaction term (FE = 11.5, padj = 6.1×10⁻⁵, n = 6), suggesting that YAP target suppression, which may participate in normal contact-inhibition quiescence, is attenuated or delayed in KRAS^G12V^ cells. In the upregulated direction, EMT (FE = 6.9, padj = 2.2×10⁻⁶, n = 12) and TGF-β signaling (FE = 5.4, padj = 0.016, n = 5) were progressively enriched in KRAS^G12V^ over time, consistent with ongoing remodeling of EC identity between day 5 and day 12.

The selective upregulation of *CCNA1* stands out against this background of global G2M suppression. It is the only direct cell cycle regulator to reach significance at either timepoint and does so at both day 5 and day 12. *CCNA2*, *CDK2*, *CCNE1*, *CDKN1A*, *CDKN2A*, and *CDKN2B* were not significantly changed at either timepoint (**Supplemental Table 1, Section A**), further supporting the view that the KRAS^G12V^ proliferative phenotype operates through selective maintenance of a subset of cell cycle machinery rather than broad cyclin/CDK dysregulation.

Collectively, the TRAP-seq data reveal a coherent two-phase temporal program: early (day 5) activation of inflammatory signaling, matrix remodeling, and a subset of Notch-associated angiogenesis genes, followed by progressive acquisition of an AVM/arterial transcriptional identity by day 12 as WT ECs quiesce. *CCNA1* was the only direct cell cycle regulator to reach significance at either timepoint and did so at both (**Supplemental Table 1, Section A**), further supporting the view that the proliferative phenotype operates through selective maintenance of a subset of cell cycle machinery rather than broad cyclin/CDK dysregulation. The delayed exit from the proliferative transcriptional program, captured by the interaction term, mirrors the phenotypic resolution of proliferation observed in the co-culture assay and provides molecular confirmation that KRAS^G12V^ cells undergo morphogenetic arrest on a delayed rather than absent schedule.

### Alpelisib and Trametinib reduce KRAS^G12V^ enhanced proliferation in co-cultures

Having established the transcriptional program driven by KRAS^G12V^ across morphogenesis, we next asked whether the signaling pathways underlying it (PI3K, MEK, and VEGFR2) are functionally required for the proliferative, migratory, and morphogenic phenotypes observed in the co-cultures. Prior to functional testing in co-culture, we confirmed target engagement in 2D culture. Alpelisib (1 µM), an FDA-approved PI3Kα-selective inhibitor, reduced pAKT and pS6 levels in KRAS^G12V^ HUVECs to near-control values (**Supplemental Figure 6A and 6C**). Trametinib (10 nM), a MEK inhibitor, similarly reduced pERK to control levels (Supplemental Figure 6B and 6D). Functional validation by EdU incorporation confirmed that Trametinib abolished KRAS^G12V^-driven proliferation, Alpelisib reduced it by approximately 50%, and Pazopanib (1 µM), a VEGFR2 inhibitor, also partially reduced proliferation (**Supplemental Figure 7A and 7B**).

We next examined the effects of these inhibitors on EC proliferation in day 5 co-cultures using the SG2M reporter and ERG staining (**Figure 8A**). Both Trametinib and Alpelisib reduced the proportion of proliferating KRAS^G12V^ cells to near-control levels, while Pazopanib produced only a partial reduction. Total ERG counts were similarly reduced with Trametinib and Alpelisib (**Figure 8B**).

**Figure 8.**
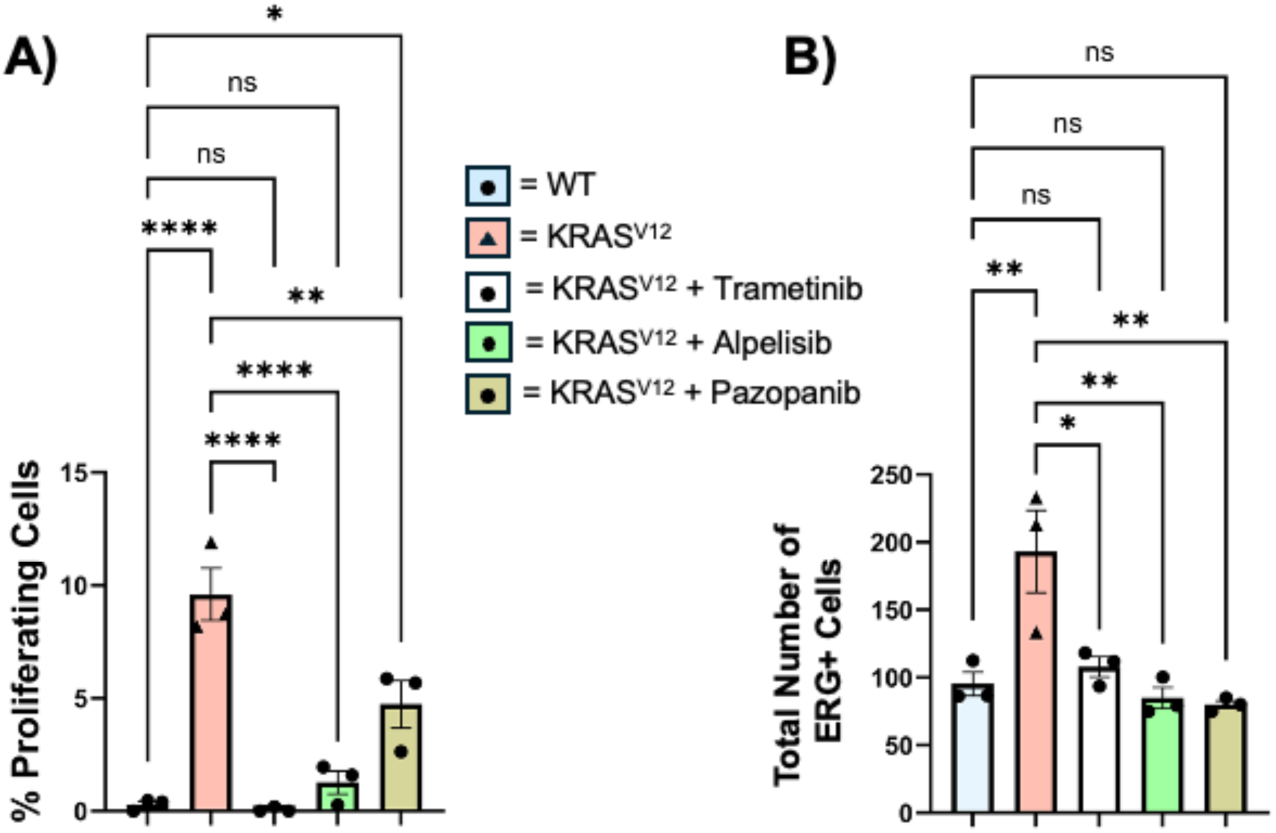
Alpelisib and Trametinib reduce proliferation to near-control levels; Pazopanib is only partially effective. A) Quantification of the proportion of proliferating cells (SG2M-positive / ERG-positive) in day 5 planar co-cultures of WT and KRAS^G12V^ HUVECs co-expressing the SG2M reporter, treated with 250 ng/mL doxycycline alone or in combination with 10 nM Trametinib, 1 μM Alpelisib, or 1 μM Pazopanib. Both Trametinib and Alpelisib reduced the proportion of proliferating KRAS^G12V^ cells to near-control levels; Pazopanib produced a partial reduction. N=4 independent experiments. One-way ANOVA, Tukey’s post-hoc test. B) Quantification of the total ERG-positive cell count per field of view for the conditions shown in A. Total EC number was reduced by Trametinib and Alpelisib treatment and partially reduced by Pazopanib. N=4 independent experiments. One-way ANOVA, Tukey’s post-hoc test, p<0.05, p<0.01, p<0.001, p<0.0001. Data represent the mean and SEM.

### Alpelisib rescues migration to near-control levels; Trametinib and Pazopanib are partially effective

The elevated *PIK3CG* expression and the established role of PI3K in EC migration [14] motivated us to test whether pharmacological inhibition of PI3K, MEK, or VEGFR2 could mitigate the enhanced migration of KRAS^G12V^ cells. We assessed migration in day 5 co-cultures by live-cell tracking in the presence of each inhibitor (**Figure 9A**). KRAS^G12V^ cells reached a mean square displacement of approximately 1,000 µm² over the 6-hour imaging period, roughly 5-fold above the ∼200 µm² observed in control cells. Alpelisib treatment reduced KRAS^G12V^ MSD to levels indistinguishable from control. Both Trametinib and Pazopanib reduced MSD by approximately 50%, to around 500 µm², but did not restore migration to control levels (**Figure 9B**). Average cell speed showed a corresponding pattern (**Figure 9C**). These results identify PI3K as the principal driver of the migratory phenotype in this system, with MAPK/ERK and VEGFR2 signaling as partial contributors.

**Figure 9.**
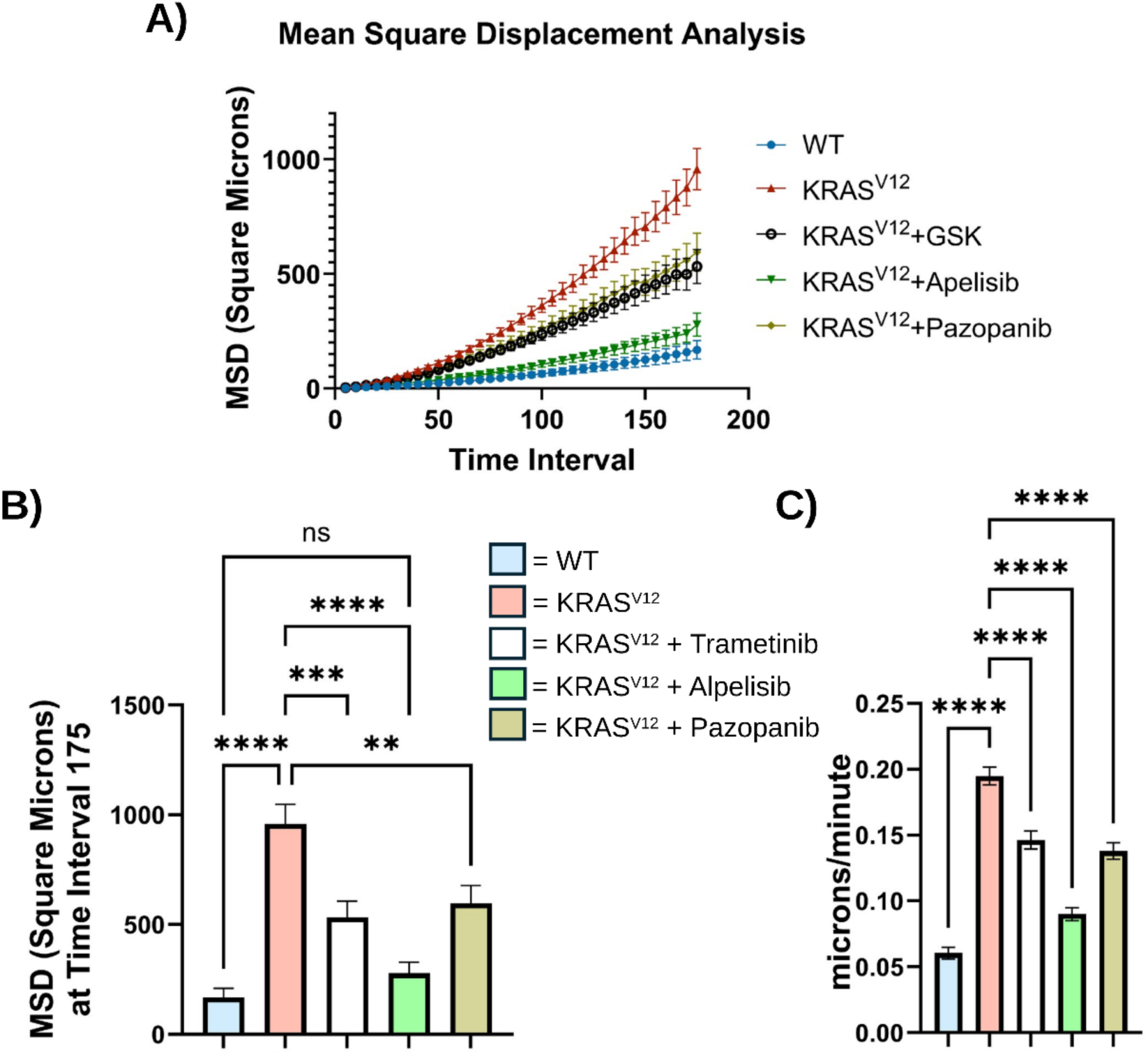
Mean square displacement and average cell speed are increased with KRAS^G12V^ and selectively reduced by Alpelisib. A) Quantification of mean square displacement (MSD) over time for 6-hour live migration tracking in day 5 planar co-cultures. Conditions: WT, KRAS^G12V^, KRAS^G12V^ + 10 nM Trametinib, KRAS^G12V^ + 1 μM Alpelisib, KRAS^G12V^ + 1 μM Pazopanib. B) Quantification of MSD at time interval 175 (6 hours). KRAS^G12V^ cells reached a significantly higher MSD than WT. Alpelisib reduced KRAS^G12V^ MSD to levels indistinguishable from WT control. Trametinib and Pazopanib each produced partial reductions but did not restore MSD to control levels. One-way ANOVA, Tukey’s post-hoc test. C) Quantification of average cell speed (μm/min) from time interval 0 to 175. The pattern of inhibitor effects on speed paralleled that observed for MSD. Tracks were generated using the mTrackJ macro in ImageJ; MSD and speed values were computed using the DiPer Excel macro. Representative of 4 independent seedings; tracks measured for WT N=127 cells, KRAS^G12V^ N=186 cells, KRAS^G12V^ + Trametinib N=143 cells, KRAS^G12V^ + Alpelisib N=133 cells, KRAS^G12V^ + Pazopanib N=159 cells. One-way ANOVA, Tukey’s post-hoc test, p<0.05, p<0.01, p<0.001, p<0.0001. Data represent the mean and SEM.

### Alpelisib restores vessel-like morphology in KRAS^G12V^ co-cultures

To determine whether these individual phenotypic improvements translated to a rescue of overall vascular architecture, we examined day 8 co-cultures treated with each inhibitor (**Figure 10A**). All three inhibitors reduced total fluorescent EC area relative to untreated KRAS^G12V^ cells (**Figure 10B**). However, only Alpelisib-treated cultures showed a meaningful restoration of vessel-like structure area as quantified by AngioTool, while Trametinib- and Pazopanib-treated cultures did not (**Figure 10C**). Representative images further illustrate that Alpelisib treatment produced thin, branched networks resembling control morphology, whereas the other two inhibitors, while reducing EC area, did not restore normal architecture. Trametinib and Pazopanib also had visually apparent negative effects on control cell morphology, while Alpelisib maintained the control network with only modest changes in tube thickness. It should be noted that inhibitors were applied to the full co-culture and effects on fibroblast-derived signals cannot be excluded (*see Discussion*).

**Figure 10.**
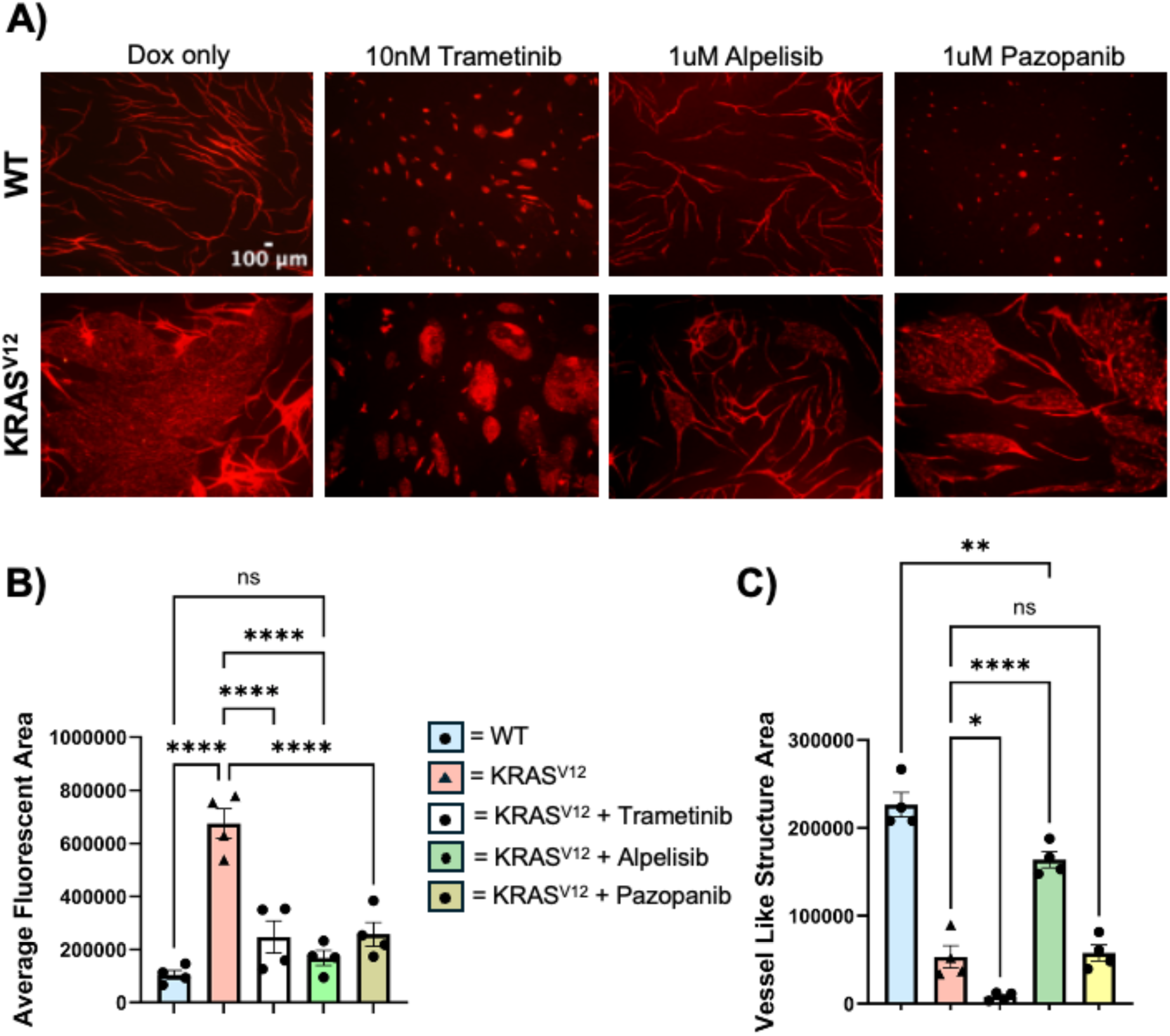
Alpelisib but not Trametinib or Pazopanib restores vessel-like morphology in KRAS^G12V^ co-cultures. A) Representative fluorescent lectin images of WT and KRAS^G12V^ planar co-cultures treated with 250 ng/mL doxycycline alone or in combination with 10 nM Trametinib, 1 μM Alpelisib, or 1 μM Pazopanib for 8 days. Alpelisib-treated KRAS^G12V^ cultures showed thin, branched networks resembling WT morphology; Trametinib- and Pazopanib-treated cultures showed reduced EC area but did not restore tubular architecture. Trametinib and Pazopanib also produced morphological changes in WT control cultures. Inhibitors were applied to the full co-culture; effects on fibroblast-derived signals cannot be excluded. Images are representative of 4 independent experiments. Scale bar = 100 μm. B) Quantification of average fluorescent area (tubes and sheets combined) by ImageJ for the indicated conditions. All three inhibitors reduced total fluorescent EC area in KRAS^G12V^ cultures. N=4 independent experiments. One-way ANOVA, Tukey’s post-hoc test. C) Quantification of vessel-like structure area by AngioTool for the indicated conditions. Only Alpelisib-treated KRAS^G12V^ cultures showed meaningful restoration of vessel-like structure area; Trametinib- and Pazopanib-treated cultures did not. N=4 independent experiments. One-way ANOVA, Tukey’s post-hoc test, p<0.05, p<0.01, p<0.001, p<0.0001. Data represent the mean and SEM.

Together, these data establish that PI3K inhibition with Alpelisib rescues all three components of the KRAS^G12V^ cellular phenotype: migration, proliferation, and morphology. MEK and VEGFR2 inhibition are sufficient to reduce proliferation and migration but insufficient to restore normal tubulogenesis.

## DISCUSSION

### The KRAS^G12V^ endothelial program

Expression of KRAS^G12V^ in primary HUVECs undergoing 3D co-culture morphogenesis produces a coherent transcriptional and phenotypic program in the endothelium: dysregulated vessel-like structure formation, enhanced migration, transient elevation of proliferation, and progressive acquisition of a transcriptional profile associated with arteriovenous malformations. Prior work from our laboratory established that activated Ras in primary endothelial cells drives a proangiogenic phenotype, with PI3K signaling identified as a key regulator of the resulting morphogenic defects [11, 12]. The present study extends that framework to the KRAS^G12V^ variant commonly identified in sporadic brain AVMs, using TRAP-seq of cells undergoing active 3D morphogenesis to provide temporal resolution of the transcriptional program [18]. Restricting KRAS expression to the EC compartment while preserving the fibroblast microenvironment captures the endothelial response in a context that retains more of the cellular complexity of developing vasculature than conventional 2D culture permits [11, 12, 18].

### Proliferation and context-dependence

Fish et al. reported that endothelial-specific KRAS expression did not alter EC number in mouse or zebrafish models, attributing the vascular phenotype to increased EC size and enhanced migration [5]. Saito et al. similarly observed only occasional Ki67 signal in lesion ECs [7]. In both models, high-efficiency delivery of mutant KRAS produces lesions in only a subset of vessels despite broad expression across the vascular beds. This incomplete penetrance is consistent with a requirement for a permissive angiogenic state for lesion initiation, as demonstrated in HHT pathway models where regional genetic mutation alone is insufficient and concurrent angiogenic stimulation is required for bAVM formation [46, 47]. A transient proliferative response may therefore accompany the angiogenic expansion that triggers lesion initiation, with cells having resolved toward quiescence by the time the nidus is accessible for measurement. EC proliferation does accompany cerebrovascular malformation in the HRAS mouse model [8], and activated Ras is well established to drive EC proliferation in primary culture [11, 12], suggesting the proliferative response is context-dependent rather than absent [5, 7, 8, 11, 12, 47, 48].

In the present study, KRAS^G12V^ cells show significantly elevated proliferation at days 3 and 5 of co-culture that resolves to near-control levels by day 12, with kinetics consistent with contact inhibition. Among direct cell cycle regulators, only CCNA1 reached significance, doing so consistently at both day 5 and day 12, suggesting selective maintenance of a subset of cell cycle machinery rather than broad cyclin/CDK dysregulation (Supplemental Table 1, Section A). Retention of G2M checkpoint genes in KRAS^G12V^ cells as WT cells downregulate them between day 5 and day 12 is consistent with a delay in morphogenetic arrest (Supplemental Table 1, Section B). The interaction contrast (Day 12 vs Day 5, KRAS vs WT) provides independent confirmation: E2F Targets (NES = −3.18), G2M Checkpoint (NES = −2.92), and MYC Targets V1 (NES = −2.86) are among the most significantly negatively enriched gene sets (all padj = 1×10⁻⁹), indicating the proliferative program active at Day 5 is largely resolved in KRAS^G12V^ cells by Day 12, even if delayed relative to WT. Proliferative gene sets are not enriched in the nidus endothelial populations of the Winkler 2022 atlas [46], suggesting the quiescent state of the mature human lesion is the endpoint of a process whose earlier proliferative phase is captured *in vitro* here.

A further observation relevant to context-dependence concerns the anatomical restriction of lesion formation in mouse models with pan-endothelial mutant Ras expression. In the iEC-HRAS^V12^ model, cerebrovascular malformations developed with high penetrance, while no vascular abnormalities were found outside the CNS [8], and Fish et al. reported brain AVMs in ibEC-Kras^G12D^ mice with retinas unaffected [5]. The basis for this CNS selectivity is not established but is consistent with the organ-specific transcriptional and functional differentiation of endothelium [49], and raises the possibility that the brain microenvironment is uniquely permissive to Ras-driven morphogenic dysregulation.

### Migration program

Enhanced migration at Day 5 is driven by two complementary programs. Within the Focal Adhesion & ECM category, ITGB3 (encoding the β3 integrin subunit) was prominently upregulated, consistent with VEGF-driven induction of αvβ3 and its established roles in Src-dependent adhesion signaling and focal adhesion remodeling during EC migration [49]. PLAU and PLAUR were co-upregulated, indicating coordinate activation of the uPA/plasmin proteolytic axis that facilitates matrix degradation and invasion [50]. Co-induction of integrin-mediated adhesion and pericellular proteolysis represents a program well-suited to 3D morphogenesis, where cells must both anchor to and remodel the surrounding matrix to advance. Among the KRAS-Direct Migration genes, PIK3CG (encoding the p110γ catalytic subunit of PI3Kγ) was significantly elevated, as were RALA, a Ras effector that promotes motility through exocyst-dependent vesicle trafficking, and RAC2, the sole Rho/Rac family member to reach significance at Day 5. All six highlighted genes remained upregulated at Day 12, indicating a sustained rather than transient migratory program across the morphogenic window.[50, 51]

MAPK/ERK and PI3K/AKT signaling both have established roles in EC migration during angiogenesis. ERK/MAPK promotes EC sprouting and survival by suppressing Rho-kinase-dependent actomyosin contractility [52], while PI3K/AKT coordinates directed motility through actin reorganization and downstream effectors, including eNOS; Akt is both necessary and sufficient for VEGF-stimulated EC migration [14, 53, 54]. Our findings differ from Fish et al., who did not detect AKT activation or PI3K-dependent migration in KRAS^G12V^ ECs. We find that KRAS^G12V^ expression elevated pAKT in HUVECs (Supplemental Figure 1), *PIK3CG* was among the most consistently upregulated KRAS-direct migration genes at both timepoints, and Alpelisib reduced migration in both 2D and 3D assays (Supplemental Figure 2; Figure 8). These data indicate PI3K is engaged downstream of KRAS^G12V^, consistent with Ras activation of PI3K across many cellular contexts [16].

### Endothelial inflammatory signature

At Day 5, SASP, Inflammatory Response, and TNFα/NF-κB signaling were the most significantly enriched upregulated gene sets, preceding the angiogenic program that expands between Day 5 and Day 12. The genes driving these enrichments included *IL1B*, *IL1A*, *IL6*, *CCL2*, *CCL5*, *CXCL1*, *CXCL3*, and *CXCL8*, with *IL1B* showing the largest fold change (log₂FC = 3.1, padj = 0.004). *IL1A* remained the most significantly upregulated cytokine at Day 12 (log₂FC = 3.8, padj = 4.6×10⁻¹⁶), and *CXCL8*, *IL1B*, and the selectins *SELE* and *SELP* also reached significance at Day 12, indicating that while the acute NF-κB pathway enrichment resolves substantially, select inflammatory mediators are sustained. Because TRAP-seq captures the translating EC transcriptome in isolation from the wild-type fibroblast compartment, this cytokine and chemokine signature originates within the KRAS^G12V^ EC itself. Active proliferation at the same timepoint, combined with the absence of OIS gene set enrichment at any contrast, argues against a classical oncogene-induced senescence interpretation, since in OIS the secretory phenotype accompanies growth arrest rather than co-occurring with active division. NF-κB activation is a well-established consequence of Ras signaling [55], and PI3K/AKT can drive NF-κB target gene induction through IKK phosphorylation [56], a connection relevant given the PI3K engagement documented in this system.

This EC-derived inflammatory program has direct parallels across independent models. Park et al. recently demonstrated that KRAS^G12V^-expressing brain ECs in an AAV-mediated bAVM mouse model produce elevated IL-6, IL-1β, TNF-α, MMP-2, and MMP-9 within the lesion nidus, and conditioned medium from KRAS^G12V^ ECs is sufficient to activate microglia and compromise tight junction expression in WT ECs [57, 58]. In vivo, this inflammatory dialogue between mutant ECs and local microglia/macrophages contributes to barrier disruption and hemorrhagic conversion and is substantially attenuated by MMP inhibition [58]. These findings extend earlier work from this laboratory showing that iEC-HRas^V12^ mice develop vascular leak and inflammatory infiltrate alongside brain lesion formation [8], indicating that inflammatory activation of the endothelium is a shared consequence of oncogenic Ras signaling regardless of isoform. Inflammatory infiltration is also documented in human bAVM tissue, with elevated IL-6 and IL-1β associated with lesion susceptibility to hemorrhage [57, 58], placing the program identified here in a clinically relevant context. Whether inflammatory activation of the endothelium contributes to the focal nature of lesion formation against a background of more widespread KRAS^G12V^ expression is not resolved by the present data but is worth direct investigation [8, 58-60].

### AVM/arterial transcriptional identity

The early migratory program connects to a second consequence of *ITGB3* upregulation that links the adhesion-proteolytic axis to arterial EC identity. Integrin β3 promotes Notch pathway activation in endothelial cells [61], and the co-upregulation of *ITGB3* with *JAG1* at Day 5, a DSL-family Notch ligand with established roles in arterial fate specification [34], suggests the early migratory program may also serve as an entry point into the Notch-dependent identity program that becomes fully established by Day 12. The Day 5 Hallmark Angiogenesis hit is small but compositionally informative: three of five genes (*JAG1*, *ROBO4*, *FLT1*) carry direct AVM or arteriovenous identity roles, while *SPHK1* and *MDK* contribute to the activated EC state through sphingosine-1-phosphate and midkine signaling, respectively [62, 63]. The selective representation of Notch components within this early hit prompted the expanded examination shown in **Figure 7**, where the same genes reappear within a substantially enlarged program.

By Day 12, KRAS^G12V^ cells maintain and progressively upregulate genes with established roles in HHT and AVM pathobiology as WT cells exit the angiogenic program. ACVRL1 (ALK1) and ENG (endoglin), the two major HHT loci whose mutations disrupt TGF-β/BMP signaling and cause hereditary vascular malformations [44, 45], are both significantly upregulated, as is BMPR2, a cooperating type II BMP receptor (Supplemental Table). The upregulation of these three genes indicates convergence on a transcriptional signature that, in genetically distinct AVM syndromes, is produced by disruption of these same components. [44, 45]

The angiopoietin/TEK node shows the same pattern. PTPRB, TEK, TIE1, and ANGPT2 are all upregulated at Day 12 through the same mechanism: WT cells reduce expression of the entire axis by roughly half as they transition to quiescence, while KRAS^G12V^ cells maintain it. PTPRB encodes VE-PTP, an endothelium-specific phosphatase that dephosphorylates TEK, VEGFR2/KDR, and VE-cadherin [62, 63]; its co-upregulation alongside TEK and KDR reflects failure to downregulate a coherent signaling node rather than a net gain of TEK activation. Co-expression of PTPRB, TEK, and ANGPT2 in nidus endothelium is a documented feature of human bAVM tissue [64], and ANGPT2 blockade prevents and rescues retinal AVM formation in a Smad4-deficient HHT mouse model [65], placing this node among candidate targets with cross-etiology relevance. [46, 64-66]

Within the Notch/arteriovenous identity program, JAG1 and HES2 are elevated at both timepoints, with NOTCH1 reaching significance only at Day 12, consistent with progressive consolidation of Notch-mediated arterial identity. Constitutively active Notch1 or Notch4 in endothelial cells is sufficient to induce bAVM in mice, with Notch-1 activation, elevated JAG1, and HES-1 documented in Notch4 mouse models and human bAVM tissue [66-68]. Conversely, endothelial Rbpj deletion causes bAVM through loss of arterial identity [69], establishing the Notch/arteriovenous identity axis as a convergence point across mechanistically distinct AVM etiologies. The KRAS^G12V^ data presented here indicate that upstream oncogenic signaling engages this same axis through selective upregulation of Notch identity output.[67-70]

KRAS^G12V^ endothelial cells also fail to acquire markers of capillary endothelial maturation that wild-type cells upregulate at Day 12. They do not upregulate FOXF2 or MFSD2A to the degree observed in WT cells undergoing maturation. FOXF2 is a forkhead transcription factor required for brain pericyte differentiation and blood-brain barrier integrity [70]; MFSD2A is a CNS endothelial-enriched lipid transporter critical for suppression of transcytosis and BBB maintenance [71, 72] that defines brain capillary identity in single-cell atlases of the human cerebrovasculature [64]. The failure to acquire these markers, rather than active loss, parallels the sustained expression of AVM identity genes documented above and points to the same underlying failure to complete the normal morphogenetic transition. [46, 71-73]

This failure of CNS maturation is not unique to our in vitro system; single-cell transcriptomic analyses of surgically resected human bAVM tissue provide a direct point of comparison. In the Winkler 2022 atlas of resected human bAVM tissue, nidus endothelial cells exhibit this same suppression of CNS capillary identity markers, including MFSD2A, alongside upregulation of pro-angiogenic and inflammatory signals [46]. The correspondence between the day 12 KRAS^G12V^ transcriptional profile and human nidus endothelium, at the gene, pathway, and ranked enrichment level (**Supplemental Table 4**), suggests that the failure of morphological resolution in co-culture reflects a stable, disease-relevant endothelial state rather than a transient response to oncogene expression.

### PI3K inhibition and translational context

The temporal trajectory of Notch signaling raises a mechanistic question directly relevant to the pharmacological findings reported in this section. *NOTCH1* increases from a sub-threshold log₂FC of 0.42 at Day 5 (padj = 0.20) to a significant log₂FC of 1.02 at Day 12 (padj < 0.001), while *JAG1* is elevated at both timepoints and continues to rise. In sprouting angiogenesis, Notch activation in stalk cells is known to induce PTEN, suppressing PI3K activity and mediating proliferative arrest [74]. *PTEN* transcript levels are unchanged here at either timepoint, arguing that transcriptional induction of PTEN is not the means by which Notch signaling could restrain PI3K activity in this system. One possibility is that oncogenic KRAS partially uncouples this Notch→PTEN→PI3K brake, with sustained RAS-GTP-driven PI3K activity overriding PTEN-dependent restraint even where PTEN protein is present [16], and progressive Notch consolidation between Day 5 and Day 12 representing a partially effective counterforce that contributes to proliferative resolution without fully normalizing PI3K output. This interpretation would predict that PI3K inhibition should be most efficacious early, when PI3K drive is highest and Notch-mediated restraint is least established. This is a consideration potentially relevant to both the *in vitro* rescue described below and the absence of rescue observed with Alpelisib in the iEC-HRasV12 model and with LY294002 in established zebrafish shunts.

Pharmacological dissection of the KRAS^G12V^ phenotype with three targeted inhibitors produced a coherent picture of pathway dependence. Alpelisib, an FDA-approved PI3Kα-selective inhibitor, restored migration to control levels in Day 5 co-cultures, reduced proliferation, and at Day 8 produced vessel-like network morphology resembling WT controls. Trametinib (MEK) and Pazopanib (VEGFR2) each reduced MSD by roughly half and suppressed proliferation comparably to Alpelisib, but neither restored normal tubulogenic architecture. The partial migration and proliferation rescue by Trametinib and Pazopanib, together with their failure to restore morphology, identifies PI3K as the principal organizer of the 3D phenotype in this system, with MAPK/ERK and VEGFR2 signaling contributing to individual components without being sufficient for morphological normalization. This is consistent with prior work from our laboratory establishing PI3K as essential for Ras-induced migratory changes while ERK activation is the principal driver of Ras-induced EC proliferation [11], and with the broader principle that PI3K/AKT coordinates directed motility and cytoskeletal organization in ways MEK/ERK activation does not fully substitute for [53].

The current data differ from Fish et al., who found that PI3K inhibition did not suppress migration in KRAS^G12V^ ECs and did not detect AKT activation downstream of KRAS^G12V^ [5]. We confirmed pAKT elevation in KRAS^G12V^ HUVECs by western blot (Supplemental Figure 1) and find Alpelisib reduces migration to control levels. The basis for this discrepancy is not fully resolved but may reflect differences in assay conditions, inhibitor concentration, or cellular context.

The in vitro rescue by Alpelisib does not translate straightforwardly to a therapeutic inference. In the iEC-HRasV12 mouse model we previously published, systemic BYL719 (Alpelisib) administered after lesion establishment not only failed to normalize the cerebrovascular phenotype but worsened lesion vessel diameter and reduced survival [8]. This does not discount the role of PI3K engagement in the EC but rather illustrates the difference between PI3K’s role in the endothelium and its broader function in the vascular microenvironment, where pericytes, microglia, astrocytes, and recruited immune cells all depend on PI3K signaling for homeostatic functions. Pan-vascular PI3K-alpha inhibition may suppress microenvironmental signals that normally contain lesion growth or support barrier integrity and maturation (for example Tie2 signaling), even while correcting endothelial morphogenesis. These types of influences are unlikely to manifest in this co-culture system, where pericyte investiture and flow are not present. The in vitro rescue identifies the pathway architecture within the endothelium; translating that finding to a therapeutic application will require approaches that concentrate on the specific endothelial target pathways within the mutant endothelium without disrupting the broader microenvironmental context on which vascular stability depends.[8]

### Limitations and future directions

Several limitations warrant consideration. The pharmacological experiments used inhibitors applied to the full co-culture, and effects on fibroblast PI3K, MEK, or VEGFR2 signaling cannot be excluded as contributors to the observed rescue. Although the TRAP-seq design restricts the transcriptional readout to the EC compartment, the inhibitor experiments do not achieve the same specificity. The lentiviral KRAS^G12V^ overexpression system in primary HUVECs, while practical, does not replicate the mosaic, low-frequency somatic mutation pattern that characterizes sporadic bAVM. Whether the transcriptional program identified here reflects what occurs in the subset of cells that go on to form lesions, rather than in the broader KRAS^G12V^-expressing population, remains unresolved by these data. The incomplete penetrance of lesion formation in mouse models with widespread endothelial Ras expression, noted above, suggests additional permissive factors govern whether a given KRAS^G12V^-expressing EC participates in lesion formation, and the relationship between those factors and the transcriptional programs characterized here remains to be defined.

TRAP-seq captures the actively translated EC transcriptome but does not report protein levels, post-translational modifications, or secreted factors. The inflammatory program described above rests on transcript-level data; while cross-model consistency with Park et al. provides partial protein-level validation for selected cytokines [57, 58], systematic characterization of the KRAS^G12V^ EC secretome and its functional consequences for surrounding cell types remains an important open question. The dialogue between mutant ECs and the microenvironment, in which mutant endothelial cells activate pericytes, astrocytes, microglia, and macrophages whose responses feedback on the lesion, was not accessible to the co-culture system used here and represents a priority for future in vivo work, particularly given the therapeutic relevance of microglial activation demonstrated by Park et al [57, 58].

The transcriptional convergence between the KRASG12V in vitro program and human bAVM nidus endothelium suggests that the co-culture system may serve as a useful screening platform for candidate interventions targeting the AVM transcriptional identity program, including the angiopoietin/TEK axis and Notch target gene activation that emerge as consistent features across model systems and human tissue. In vivo validation in genetically defined models, where the full neurovascular microenvironment is preserved, will be required to determine which of these candidates has therapeutic relevance beyond the in vitro endothelial context.

## CONFLICT OF INTEREST STATEMENT

The authors have no conflicts of interest to disclose.

## FIGURE LEGENDS

**Supplemental Figure 1. KRASG12V expression increases levels of active AKT, S6, and ERK in the absence of growth factors.**

A) Representative blot of Pan-Ras (Myc-tagged Ras and endogenous Ras), GAPDH, and ERK1/2 from HUVECs serum-starved and induced with 50 ng/mL doxycycline for 24 hours prior to lysis. Lanes 1 and 2 represent two replicates of one experiment for the WT lysate; lanes 3 and 4 represent two replicates of one experiment for the KRAS^G12V^ lysate.

B) Quantification of the blot shown in A (N=6). KRAS^G12V^ expression level is calculated as the ratio of Myc-tagged Ras to endogenous Ras as detected by the Pan-Ras antibody, normalized to GAPDH. KRAS^G12V^ expression is approximately 3.5-fold above endogenous KRAS.

C) Representative blots of pAKT, total AKT, pERK1/2, total ERK2, pS6, and total S6 from HUVECs serum-starved and induced with 0, 50, or 250 ng/mL doxycycline for 24 hours prior to lysis. Images are representative of 3–5 independently seeded experiments.

D) Quantification of the blots shown in C. pAKT, pS6, and pERK1/2 levels are shown relative to their total protein and normalized to the WT at 0 ng/mL doxycycline. N=3–5 independently seeded experiments. P-values are determined by Student’s T-test, p<0.05, p<0.01, p<0.001, p<0.0001. Data represent the mean and SEM.

**Supplemental Figure 2. Functions important for angiogenesis are disrupted in 2D culture of KRAS^G12V^ HUVECs.**

A) Quantification of 2D proliferation in HUVECs induced with 50 ng/mL doxycycline and serum-starved for 36 hours before incubation with EdU for 4 hours, followed by fixation and staining for EdU and DAPI. The ratio of EdU-positive to DAPI-stained cells is shown. KRAS^G12V^ cells showed significantly higher EdU incorporation than control under serum-free conditions; no difference was observed in complete growth medium. N=4–6 independent experiments. Student’s T-test.

B) Quantification of transwell migration. HUVECs serum-starved and induced with 50 ng/mL doxycycline were allowed to migrate across a transwell membrane for 4 hours prior to fixation and DAPI staining. KRAS^G12V^ cells showed significantly greater numbers of migrated cells compared to control. N=6 independent experiments. Student’s T-test.

C) Survival analysis in HUVECs cultured in serum-free M199 or complete growth medium (ECGM) for 72 hours with 50 ng/mL doxycycline prior to lysis. The percent of cleaved caspase-3 relative to total caspase-3 is shown. KRAS^G12V^ cells had significantly lower cleaved caspase-3 than control under serum-free conditions, consistent with a survival advantage. No difference was observed in complete growth medium. N=5 independent experiments. P-values are determined by Student’s T-test, p<0.05, p<0.01, p<0.001, p<0.0001. Data represent the mean and SEM.

**Supplemental Figure 3. Average cell size is increased in KRAS^G12V^ HUVECs.**

A) Representative images of WT and KRAS^G12V^ cells in serum-free medium (SFM) and complete growth medium (CGM) following treatment with 50 ng/mL doxycycline for 24 hours. Cells were stained with fluorescent lectin (cytoplasm) and ERG (nuclei). Images are representative of 4 independent experiments. Scale bar = 100 μm; scale bar applies to all images.

B) Quantification of Feret’s diameter, determined by ImageJ analysis of lectin-stained images. HUVECs were seeded, allowed to attach, and then placed in SFM or CGM with 50 ng/mL doxycycline for 24 hours prior to fixation. KRASG12V cells had significantly greater Feret’s diameter in serum-free conditions; the difference in complete growth medium did not reach significance. N=4 independent experiments.

C) Quantification of average area per cell for the same conditions as B. Average area per cell was calculated by dividing total lectin area by total ERG-positive cell count per field of view. KRASG12V cells had significantly greater area per cell in both conditions. N=4 independent experiments. P-values are determined by Student’s T-test, *p<0.05, **p<0.01, ***p<0.001,

****p<0.0001. Data represent the mean and SEM.

**Supplemental Figure 4. TRAP-seq quality control: variance explained by principal components and inter-vendor expression concordance.**

A) Percentage of variance explained by each of the top four principal components, computed across all 16 TRAP-seq samples. PC1 (45.8%) reflects donor lot of origin; PC2 (21.3%) separates samples by genotype and timepoint.

B, C) Scatter plots comparing mean normalized expression per gene between PromoCell and Lonza donor lots at Day 5 (B) and Day 12 (C). The dashed line indicates y = x; the red line shows the linear regression fit. Pearson r = 0.960 at Day 5 and r = 0.967 at Day 12, confirming high inter-vendor concordance.

**Supplemental Figure 5. Expression of Notch pathway genes across eight functional categories in KRAS^G12V^ HUVECs.**

Heatmap of VST-normalized expression Z-scores for 47 Notch-associated genes reaching significance in at least one contrast (KRAS^G12V^ vs. WT at day 5, day 12, or the day 12 vs. day 5 interaction term; adjusted p < 0.05). Values represent group means across four independent HUVEC donors (N = 4) computed from variance-stabilizing transformed counts and Z-scored per gene across the four group means (WT day 5, WT day 12, KRAS^G12V^ day 5, KRAS^G12V^ day 12); color scale is clipped at ±1.5. Genes are organized into eight functional categories as defined in the Methods (color-coded sidebar); within each category, genes are sorted by descending log₂ fold change at day 12. Significance annotations to the right of the heatmap indicate the contrast in which each gene reached significance: D5 (KRAS^G12V^ vs. WT at day 5), D12 (KRAS^G12V^ vs. WT at day 12), and Inter (day 12 vs. day 5 interaction term). Star color indicates direction of effect: red denotes higher expression in KRAS^G12V^ relative to WT; blue denotes lower expression. p < 0.05, p < 0.01, p < 0.001, p < 0.0001 (DESeq2, Benjamini–Hochberg adjusted). Two genes present in both the Receptors and AVM/Arteriovenous Identity categories (NOTCH1, NOTCH4) are shown once under Receptors.

**Supplemental Figure 6. Protein-level validation of target engagement for Trametinib and Alpelisib.**

A) Representative blots of pS6, total S6, pAKT, and total AKT from lysates of WT, KRAS^G12V^, and KRAS^G12V^ treated with 1 μM Alpelisib for 24 hours in endothelial complete growth medium. Representative of 5 independent experiments.

B) Representative blots of pERK1/2 and total ERK1/2 from lysates of WT, KRAS^G12V^, and KRAS^G12V^ treated with 10 nM Trametinib for 24 hours in endothelial complete growth medium. Representative of 4 independent experiments.

C) Quantification of pAKT and pS6 from the blots shown in A, normalized to their respective total protein levels and to WT. Alpelisib reduced both pAKT and pS6 in KRAS^G12V^ cells to near-control levels. N=5 independent experiments.

D) Quantification of pERK1/2 from the blots shown in B, normalized to total ERK1/2 and to WT. Trametinib reduced pERK1/2 in KRAS^G12V^ cells to near-control levels. N=4 independent experiments. P-values are determined by Student’s T-test, p<0.05, p<0.01, p<0.001, p<0.0001. Data represent the mean and SEM.

**Supplemental Figure 7. Functional validation of Trametinib, Alpelisib, and Pazopanib on KRAS^G12V^-driven proliferation in 2D culture.**

A) Representative images of WT and KRAS^G12V^ HUVECs treated with 250 ng/mL doxycycline alone or in combination with 10 nM Trametinib, 1 μM Alpelisib, or 1 μM Pazopanib for 24 hours in endothelial complete growth medium, stained for EdU and DAPI. Images are representative of 4 independent experiments. Scale bar = 100 μm.

B) Quantification of the percent of proliferating cells (EdU-positive / DAPI-positive) for the conditions shown in A. Trametinib abolished KRAS^G12V^-driven proliferation; Alpelisib reduced it by approximately 50%; Pazopanib produced a partial reduction. N=4 independent experiments. One-way ANOVA, Tukey’s post-hoc test, p<0.05, p<0.01, p<0.001, p<0.0001. Data represent the mean and SEM.

## Notes

### Competing Interest Statement

The authors have declared no competing interest.

### Funding

- National Heart Lung and Blood Institute: R01 HL151679
- National Institute on Aging: U01 AG072464
- David E. Bryant Trust

