## Supplementary figures and images for "Endothelial KRAS^G12V^ signaling drives aberrant morphogenesis and establishes an AVM transcriptional identity in primary human endothelial cells"

### Supplemental Figures

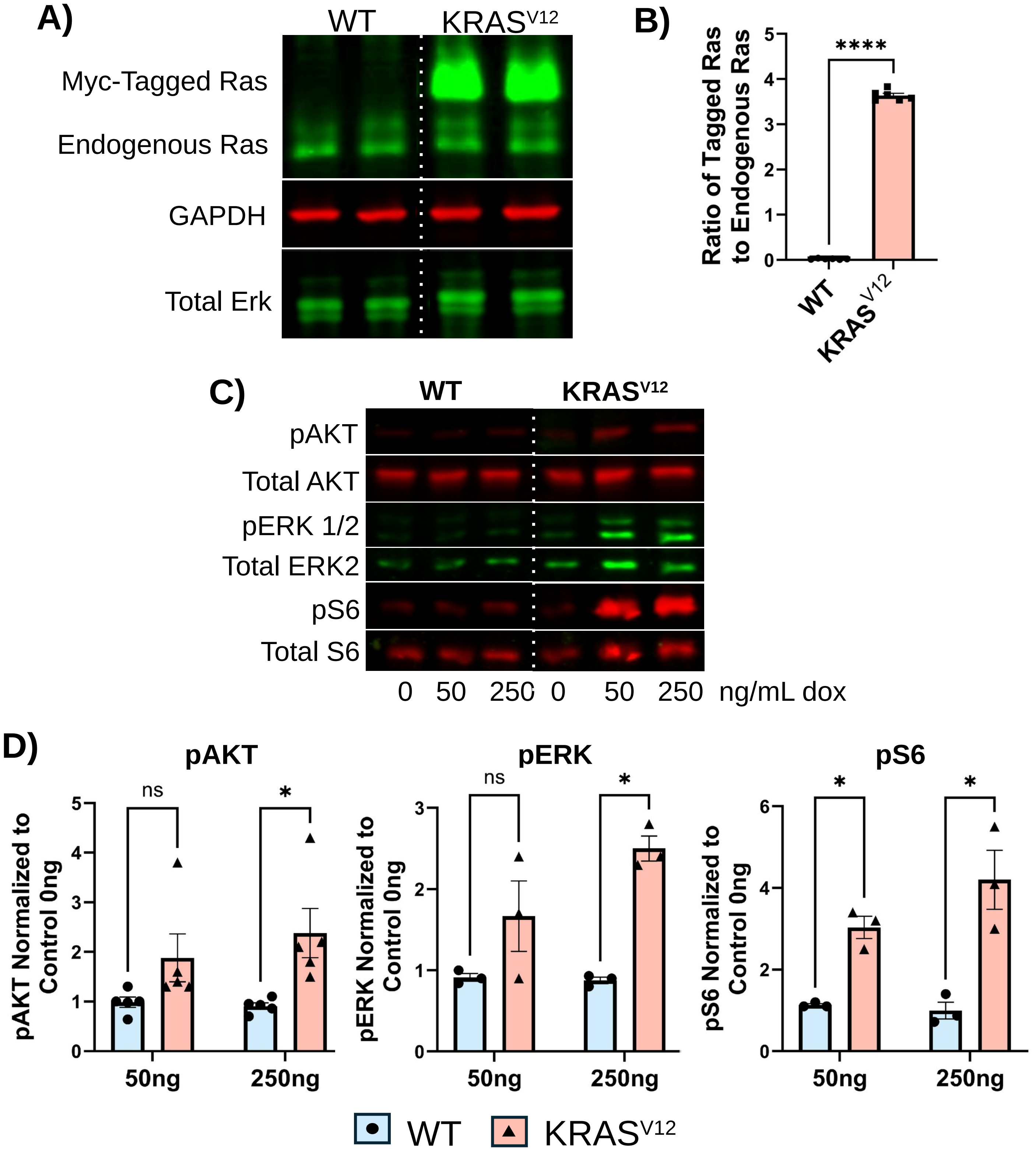

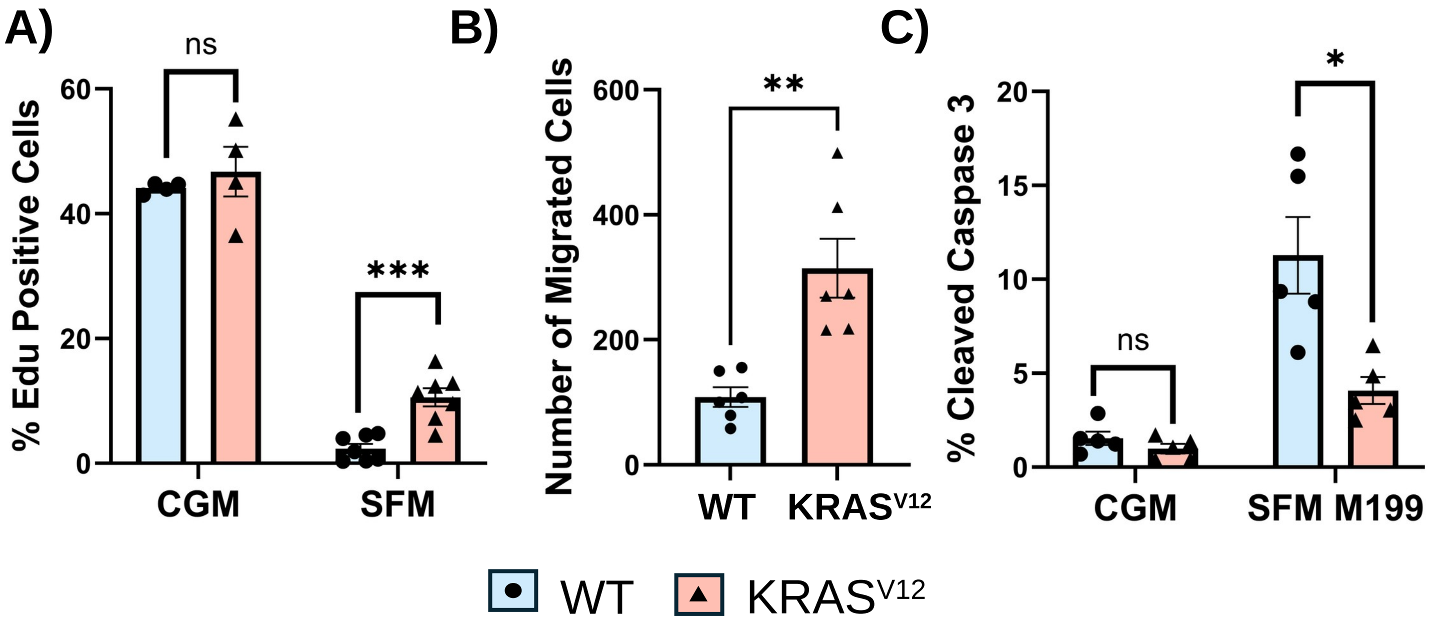

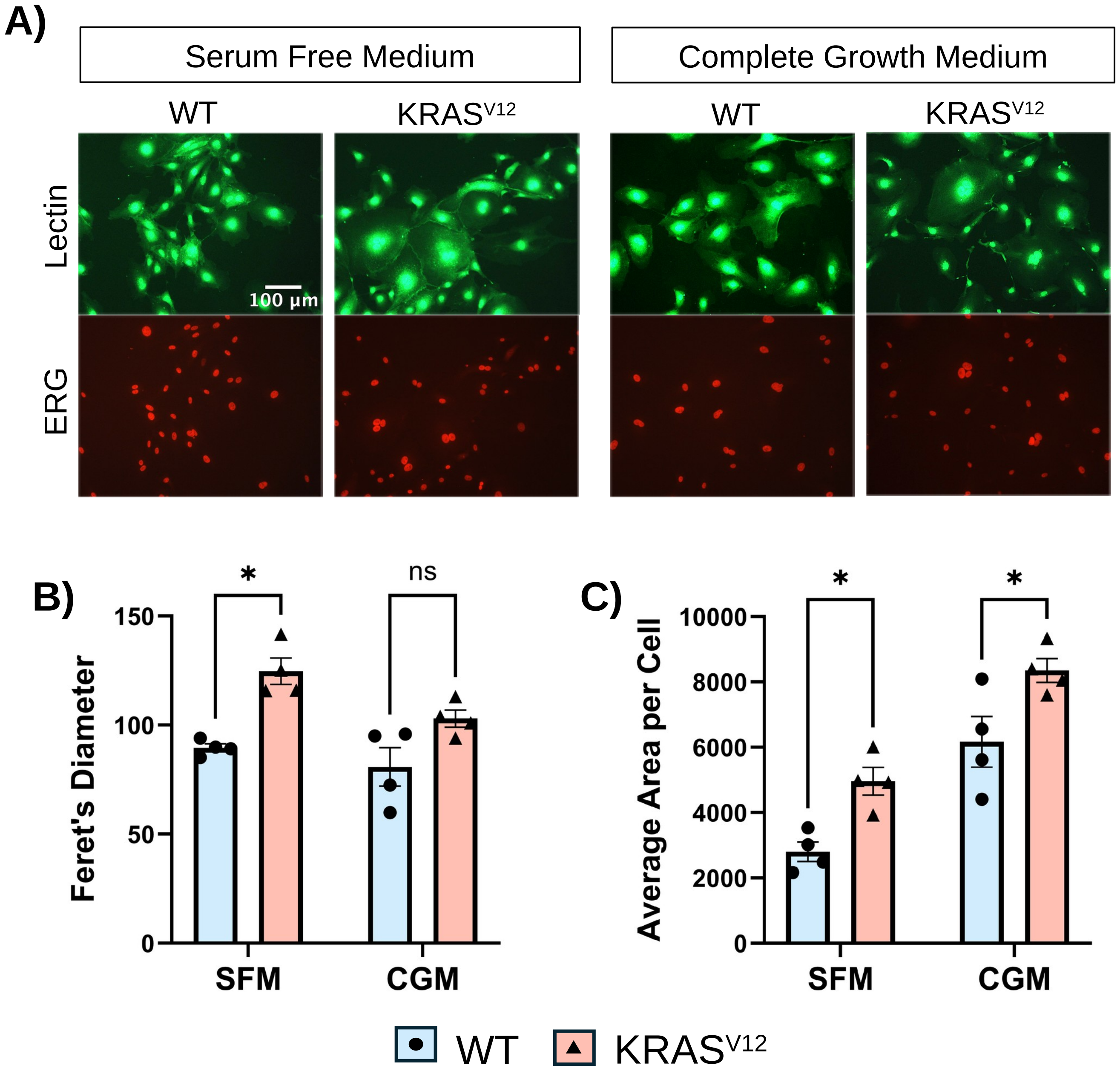

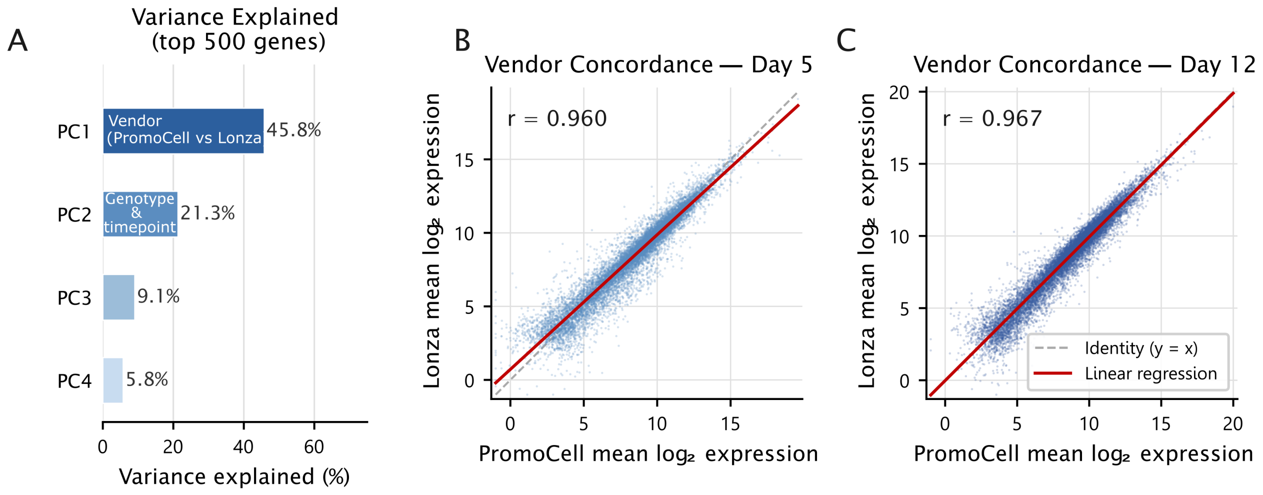

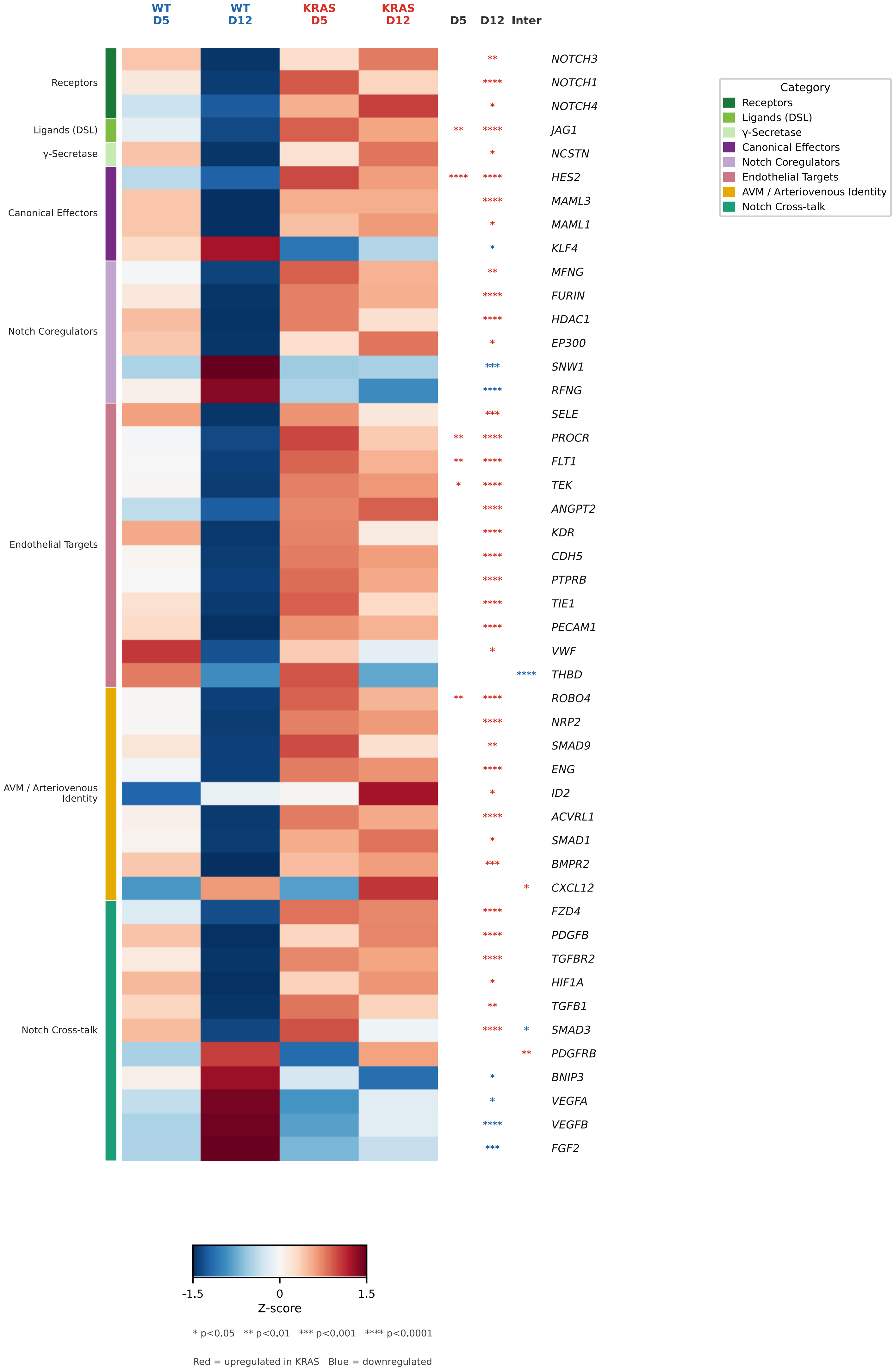

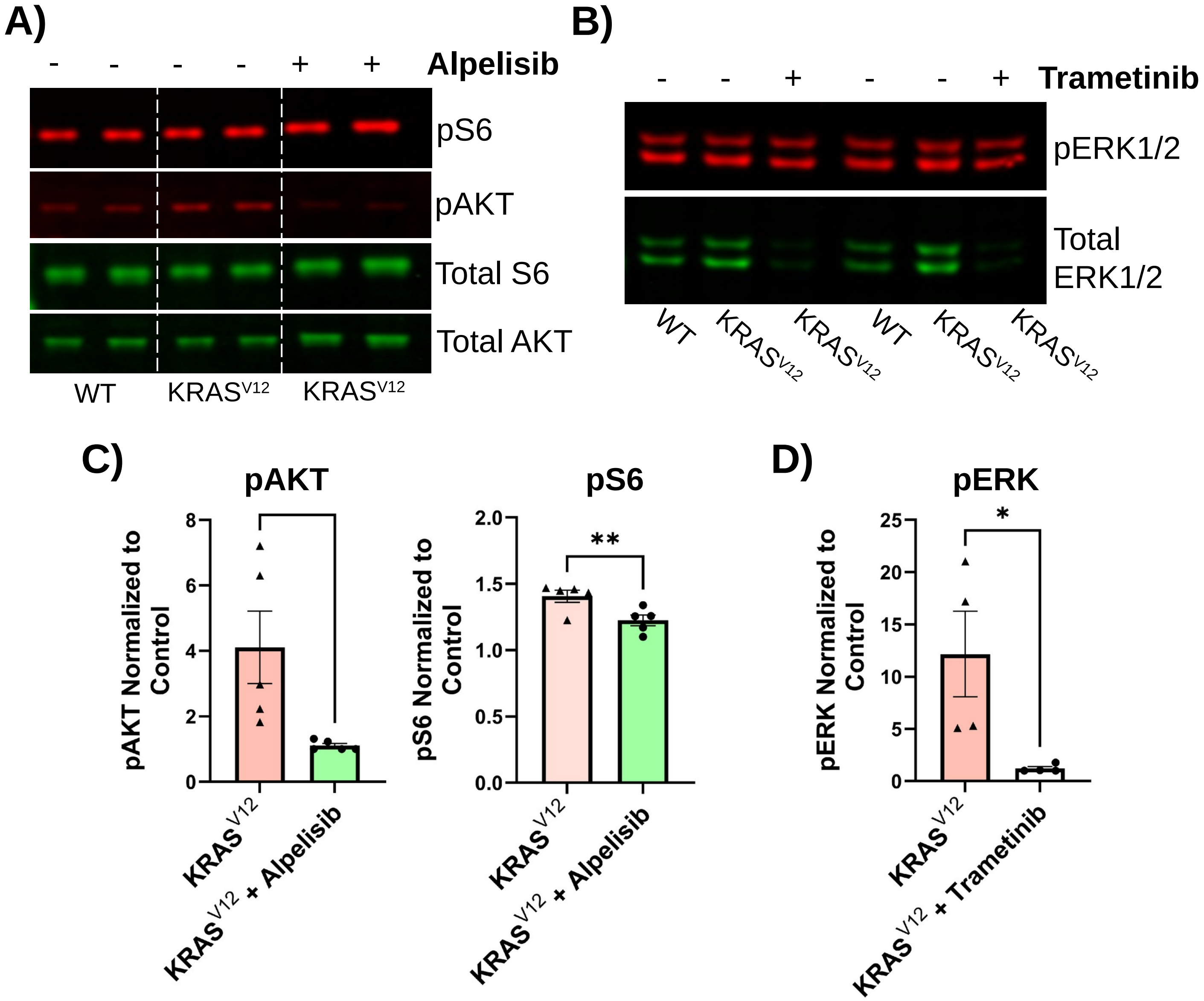

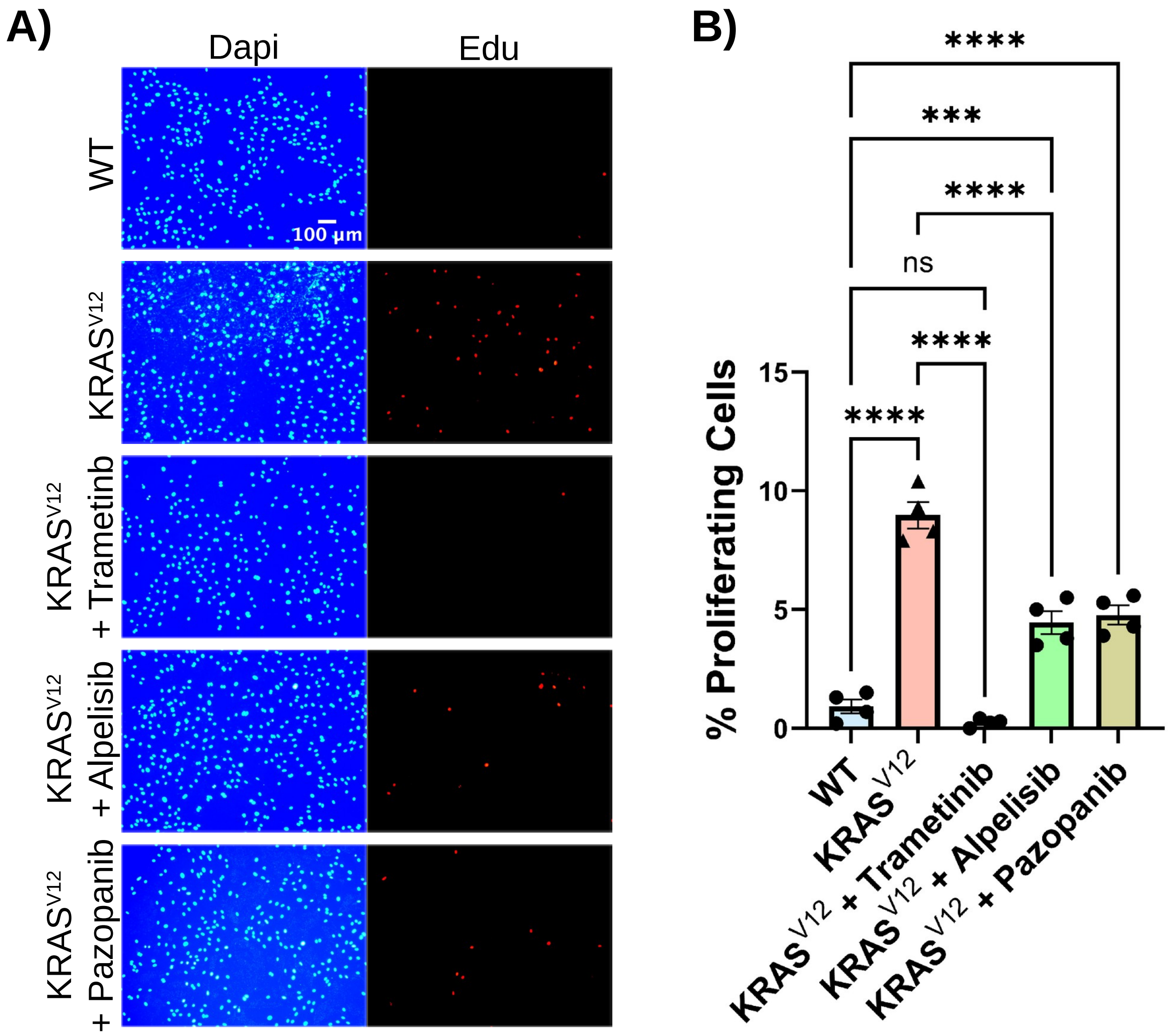
